# A dynamic equilibrium between TTP and CPEB4 controls mRNA stability and inflammation resolution

**DOI:** 10.1101/2021.03.11.434803

**Authors:** Clara Suñer, Annarita Sibilio, Judit Martín, Chiara Lara Castellazzi, Oscar Reina, Ivan Dotu, Adrià Caballé, Elisa Rivas, Vittorio Calderone, Juana Díez, Angel R. Nebreda, Raúl Méndez

## Abstract

Temporal control of inflammation is critical to avoid pathological developments, and is largely defined through the differential stabilities of mRNAs. While TTP-directed mRNA deadenylation is known to destabilize ARE-containing mRNAs, this mechanism alone cannot explain the variety of mRNA expression kinetics observed during inflammation resolution. Here we show that inflammation resolution requires CPEB4 expression, *in vitro* and *in vivo*. Our results identify that CPEB4-directed polyadenylation and TTP-mediated deadenylation compete during the resolutive phase of the LPS response to uncouple the degradation of pro-inflammatory mRNAs from the sustained expression of anti-inflammatory mRNAs. The outcome of this equilibrium is quantitatively defined by the relative number of CPEs and AREs in each mRNA, and further shaped by the coordinated regulation by the MAPK signalling pathway of the levels and activities of their trans-acting factors, CPEB4 and TTP. Altogether, we describe a temporal- and transcript-specific regulatory network controlling the extent of the inflammatory response.

## INTRODUCTION

As part of the innate immune system, macrophages sense infectious pathogens and orchestrate inflammatory and antimicrobial immune responses. These immune reactions require tight temporal regulation of multiple pro- and anti-inflammatory factors, connected by negative feedback loops, which ultimately ensure inflammation resolution. Alterations of these dynamic and coordinately regulated gene expression programs cause pathological inflammation and lead to diseases such as auto-immune disorders and cancer (Carpenter et al., 2014). While changes in the rate of gene transcription are important for the initial inflammatory response, its duration and strength are determined mainly by the rate of mRNA decay (Rabani et al., 2011).

AU-Rich Elements (AREs) are key cis-acting elements that regulate mRNA deadenylation and the stability of transcripts involved in inflammation (Spasic et al., 2012). The role of the ARE-binding protein Tristetraprolin (TTP) has been widely characterized in macrophages stimulated by bacterial lipopolysaccharide (LPS) (Carpenter et al., 2014). During the late LPS response, TTP-mediated mRNA decay limits the expression of inflammatory genes, thereby establishing a post-transcriptional negative feedback loop that promotes the resolution of inflammation (Anderson, 2010; Spasic et al., 2012). However, early in the LPS response, TTP activity is counterbalanced by another ARE-binding protein, Hu-antigen R (HuR), which stabilizes its mRNA targets by competing with TTP for ARE occupancy (Tiedje et al., 2012). The competitive binding equilibrium between TTP and HuR is post-translationally regulated by the mitogen-activated protein kinase (MAPK) signalling pathways, whose activation is induced by LPS through Toll-like receptor 4 (TLR4) (Arthur and Ley, 2013; O’Neil et al., 2018).

Despite extensive work, ARE-mediated deadenylation and mRNA decay do not explain the variety of temporal expression patterns and destabilization kinetics needed to orchestrate inflammation resolution (Sedlyarov et al., 2016). Thus, additional mechanisms would be required to define the temporal expression patterns of pro- and anti-inflammatory genes. In early development, the cytoplasmic regulation of poly(A) tail length is not a unidirectional event but rather the reflection of a dynamic equilibrium between deadenylation and polyadenylation by ARE-binding proteins and cytoplasmic polyadenylation element binding proteins (CPEBs) (Belloc and Mendez, 2008; Nousch et al., 2019; Pique et al., 2008). CPEBs bind to the cytoplasmic polyadenylation elements (CPEs) present in the 3’ UTR of some mRNAs and promote translation of these mRNAs by favouring the elongation of their poly(A) tails (Ivshina et al., 2014; Weill et al., 2012). The activities exerted by CPEBs are quantitatively defined by the number and position of CPEs present in their target transcripts, thus determining the polyadenylation kinetics and the transcript-specific temporal patterns of translation.

The global contribution of CPEBs to the regulation of mRNA expression during inflammatory processes has not been addressed. In this work, we show that myeloid CPEB4 is needed for the resolution of the LPS-triggered inflammatory response. We further show that the levels and activities of CPEB4 and TTP are sequentially regulated by LPS-induced MAPK signalling. In turn, the combination of CPEs and AREs in their target-mRNAs generate transcript-specific decay rates. These two opposing mechanisms allow destabilization of inflammatory transcripts while maintaining the expression of the negative feedback loops required for efficient inflammation resolution.

## RESULTS

### Inflammation resolution is impaired in *Cpeb4*KO macrophages

To determine a potential contribution of CPEBs to the regulation of inflammatory responses, we interrogated the expression of *Cpeb*-encoding mRNAs during the course of a systemic inflammatory response in sepsis patients. In two independent GEO datasets, we observed that *Cpeb4* mRNA levels were significantly upregulated in blood, whereas *Cpeb1-3* mRNAs did not present major expression changes **(Figure 1A)**. Monocytes/macrophages, which are enriched during sepsis, are the immune cell populations expressing higher *Cpeb4* mRNA levels **(Figure S1A, S1B)**. Moreover, deconvolution analysis suggested that myeloid cells express higher levels of *Cpeb4* mRNA during sepsis **(Figure S1C, S1D).** To further explore the significance of this correlation, we generated a myeloid-specific *Cpeb4* knockout mouse model (*Cpeb4*MKO) and challenged these mice with an intraperitoneal dose of LPS, the main membrane component of gram-negative bacteria. Upon LPS-induced endotoxic shock, *Cpeb4*MKO mice displayed lower survival rates than the wild-type controls **(Figure 1B)**. *Cpeb4*MKO animals also presented splenomegaly **(Figure 1C)** and increased splenic levels of cytokines *Il6*, *Tnf*, and *Il1a* **(Figure 1D)**. These results link CPEB4 ablation in myeloid cells to an exacerbated inflammatory response, which impairs survival from sepsis.

**Figure 1.**
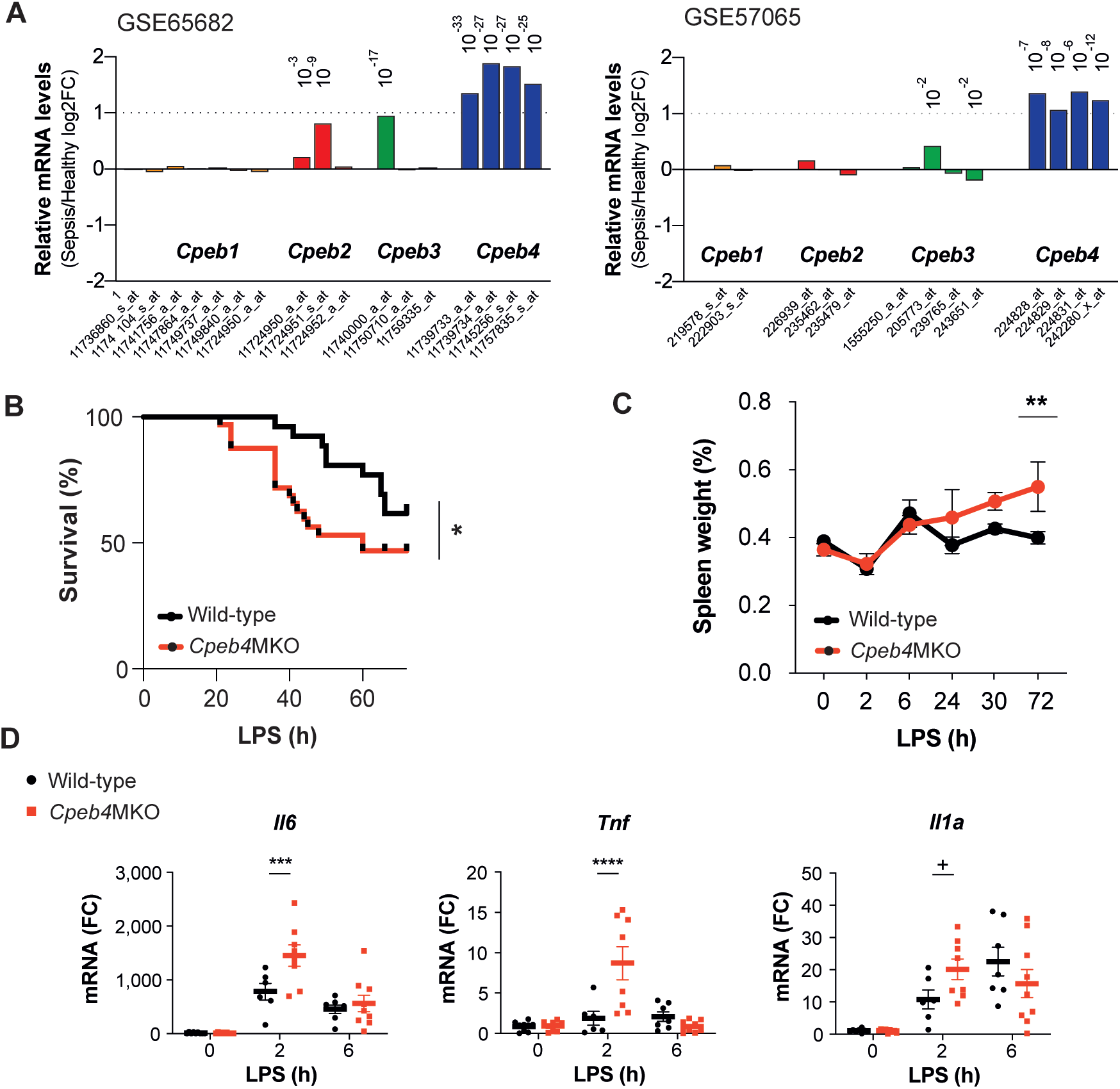
CPEB4 expression in myeloid cells is required for survival from sepsis. **(A)** Differential expression of *Cpeb* mRNAs in the blood of sepsis patients/healthy individuals. Statistics: limma moderated t-test. Pvadj (Benjamini-Hochberg) is shown. **B-D** Wild-type and myeloid-specific *Cpeb4*KO mice (*Cpeb4*MKO) were injected with an i.p. dose of LPS. **(B)** Kaplan-Meier survival curves. Results represent three independent experiments (n>7/group/experiment). Statistics: Likelihood-ratio test-pv. **(C)** Spleen weights normalized to total animal weight. Statistics: Two-way ANOVA. **(D)** Splenic total mRNA was measured by RT-qPCR and referred to *Tbp* (n>5 animals/condition). Statistics: Multiple t-test. **(C, D)** Data are represented as mean ± SD.

To further characterize CPEB4 function during the LPS response, we stimulated Bone Marrow-Derived Macrophages (BMDMs) with LPS. Time-course analysis showed that *Cpeb4* mRNA was transiently upregulated, peaking between 3 and 6 h after LPS stimulation, while the other *Cpeb* mRNAs remained virtually unaffected **(Figure 2A)**. This peak of *Cpeb4* mRNA expression was followed by CPEB4 protein accumulation **(Figure 2B)**. The CPEB4 protein was detected as a doublet with the slow migrating band corresponding to the hyperphosphorylated and active form (Guillén-Boixet, 2016) **(Figure 2C)**. Consistent with the observation that phosphorylated CPEB4 accumulated during the late phase of the LPS response, comparative transcriptomic analysis between wild-type and *Cpeb4*KO BMDMs showed that most differences appeared 6-9 h after LPS stimulation, during the resolution phase of the inflammatory response **(Figure 2D, Table S1)**. The transcriptomic profile of *Cpeb4*KO BMDMs included increases in hypoxia, glycolysis and mTOR pathways, which have been linked to pro-inflammatory macrophage polarization during sepsis (Shalova et al., 2015) **(Figure S2A)**. We confirmed that HIF1α levels were increased, both at protein and mRNA levels, after 9 h of LPS treatment **(Figure 2E, 2F)**. Conversely, anti-inflammatory transcripts like *Il10* mRNA were reduced in the *Cpeb4*KO BMDMs **(Figure 2G, Figure S2B).** All together, these results suggested that the absence of myeloid CPEB4 disrupts the homeostatic switch towards inflammation resolution.

**Figure 2.**
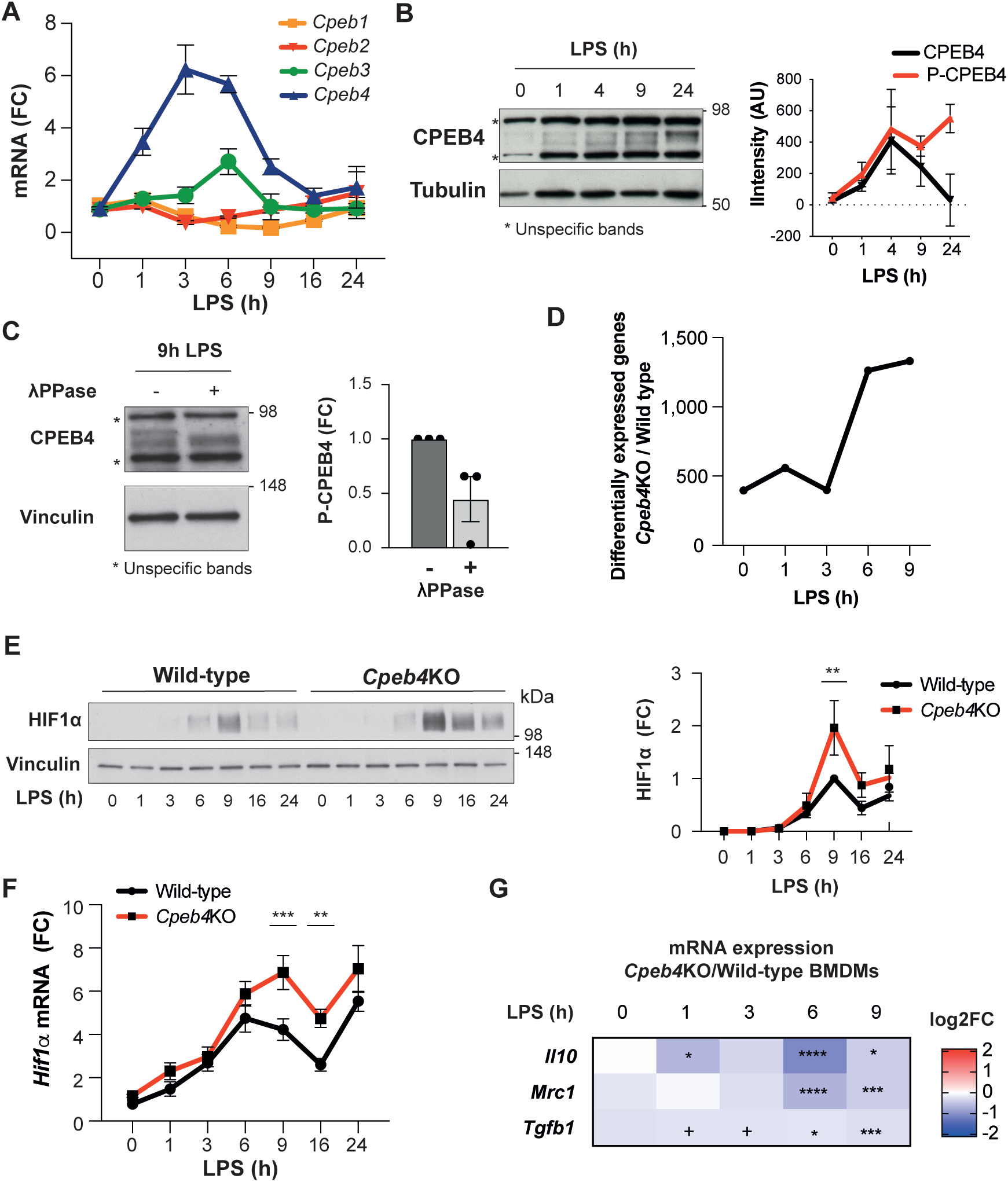
Inflammation resolution is impaired in *Cpeb4*KO macrophages. **A-C** LPS-stimulated wild-type BMDMs. **(A)** *Cpeb1-4* levels were measured by RT-qPCR (n=6). **(B)** (Left) CPEB4 immunoblot, α-Tubulin served as loading control. (Right) Quantification (n=3). **(C)** (Left) CPEB4 immunoblot in protein extracts treated with λPhosphatase when indicated. (Right) Quantification of P-CPEB4 signal (n=3). **D-G** LPS-stimulated wild-type and *Cpeb4*KO BMDMs. **(D)** Number of differentially expressed genes (pv<0.01) between genotypes. mRNA levels were quantified by RNASeq (n=4). Statistics: Deseq2 R package. **(E)** (Left) HIF1α immunoblot, vinculin served as loading control. (Right) Quantification, signal intensity was normalized to wild-type 9 h LPS (n=3). **(F)** *Hif1a* levels measured by RT-qPCR (n=6). **(G)** Differential mRNA expression measured by RNASeq (n=4). Statistics: Deseq2 R package. **(A, F)** *Tbp* was used to normalize. **(B-C, E-F)** Data are represented as mean ± SEM. **(E-F)** Statistics: Two-way ANOVA. **(D, G)** See also Table S1.

### The p38*α*-HuR-TTP axis regulates *Cpeb4* mRNA stability

We next addressed how CPEB4 expression was regulated in BMDMs treated with LPS. Since the peak of *Cpeb4* mRNA followed similar kinetics to pro-inflammatory ARE-containing mRNAs like *Il6* or *Il1a* **(Figure S3A**), we searched *Cpeb4* 3’ untranslated region (3’-UTR) for AREs and found 17 repeats of the AUUUA pentanucleotide **(Figure S3B**). ARE-containing transcripts are stabilized through HuR binding during the early phase of the LPS response and destabilized at later times by TTP binding. The switch between these two ARE-binding proteins is regulated by LPS-activated p38α MAPK (Tiedje et al., 2012). To determine whether *Cpeb4* expression was controlled by the p38*α*-HuR-TTP axis, we used BMDMs from myeloid-specific p38α KO mice (*p38α*MKO) (Youssif et al., 2018) and found that *Cpeb4* expression levels were indeed reduced in LPS-treated *p38α*MKO macrophages **(Figure 3A)**. This observation was confirmed by treating wild-type BMDMs with the p38*α* inhibitor PH797804 **(Figure 3B)**. To further define whether p38α regulates *Cpeb4* expression at the transcriptional or post-transcriptional levels, we inhibited transcription in LPS-treated wild-type and *p38α*MKO BMDMs, and found that *Cpeb4* mRNA levels decayed significantly faster in the latter **(Figure 3C)**. Next, we assessed whether the stabilization of *Cpeb4* mRNA by p38α was mediated through the differential binding of HuR and TTP. By measuring *Cpeb4* mRNA co-immunoprecipitated with HuR from wild-type and *p38α*MKO BMDMs, we observed that HuR binding to *Cpeb4* mRNA was strongly enriched upon LPS treatment in a *p38α***-**dependent manner. Similar results were observed in *Tnf* mRNA as a control **(Figure 3D, Figure S3C)**. To determine TTP binding to *Cpeb4* mRNA, we took advantage of a recent study reporting TTP immunoprecipitation (iCLIP) followed by genome-wide analysis of its associated mRNAs in LPS-stimulated BMDMs (Sedlyarov et al., 2016). Analysis of these datasets showed binding of TTP to the *Cpeb4* 3’-UTR 6 h after LPS treatment **(Figure S3D)**, as well as significantly reduced decay rates of both *Cpeb4* and *Tnf* mRNAs in TTPMKO macrophages **(Figure 3E)**. To further analyse the contribution of p38α signalling to *Cpeb4* mRNA stability, we used human osteosarcoma (U2OS) cells expressing the p38 MAPK activator MKK6 under the control of the TET-ON promoter(Trempolec et al., 2017). MKK6-induced p38α activation was sufficient to increase *Cpeb4* mRNA levels in these cells, and this effect was reversed by treatment with p38α chemical inhibitors **(Figure 3F)** or by shRNA-mediated depletion of HuR **(Figure 3G, Figure S3E, S3F, S3G)**. All together, these results indicate that *Cpeb4* mRNA is stabilized by HuR and destabilized by TTP, and that the competitive binding of these two ARE-binding proteins to *Cpeb4* mRNA is regulated by LPS-driven p38α activation.

**Figure 3.**
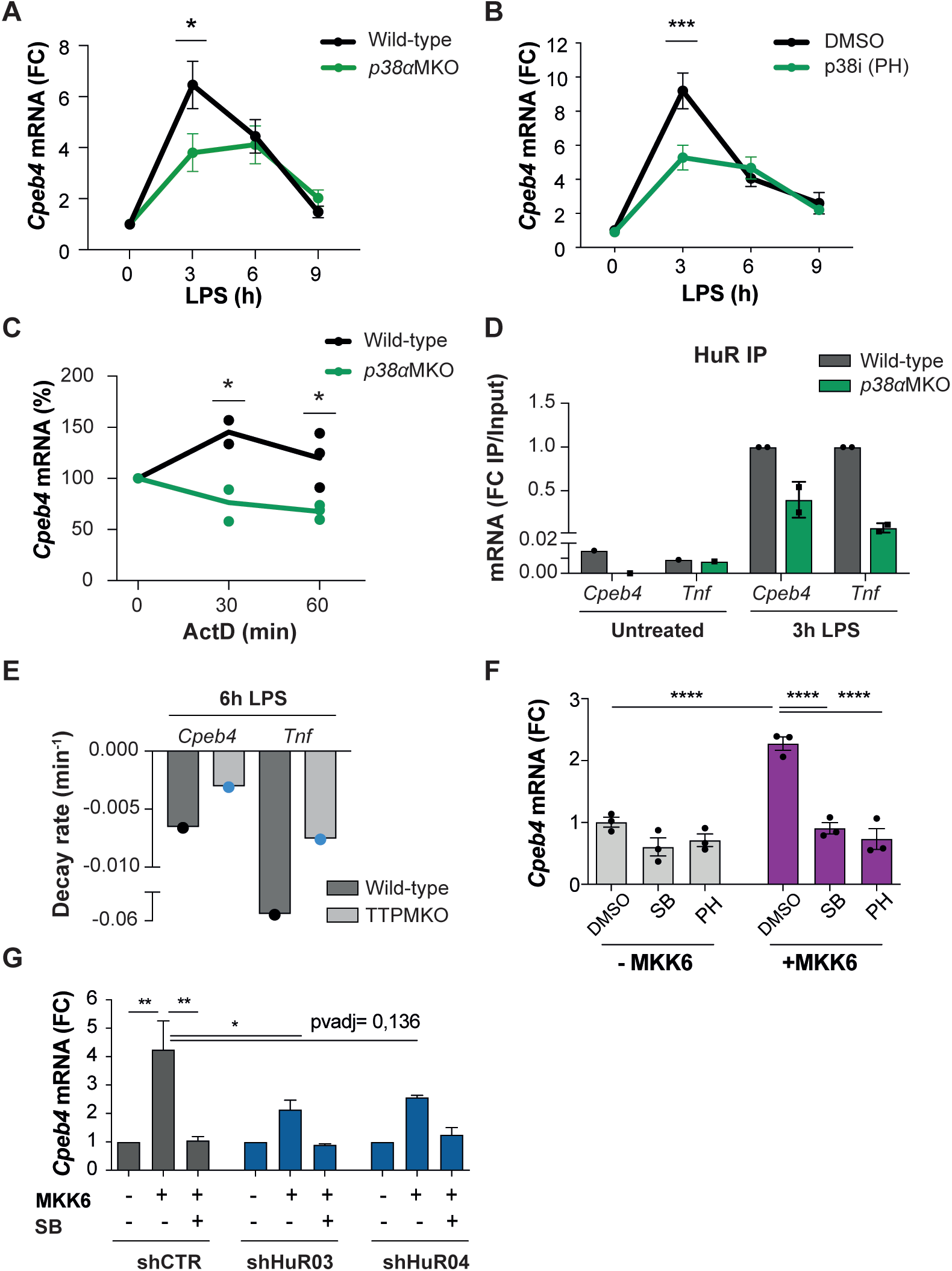
The p38*α*-HuR-TTP axis regulates *Cpeb4* mRNA stability. **(A)** *Cpeb4* levels in wild-type and *p38α*MKO BMDMs stimulated with LPS (n=3). **(B)** *Cpeb4* levels in LPS-stimulated BMDMs treated with the p38 inhibitor PH or DMSO as control (n=4). **(C)** *Cpeb4* mRNA stability was measured by treating with actinomycin D (ActD) wild-type and *p38α*MKO BMDMs stimulated with LPS for 1 h (n*≥*2). Statistics: Multiple t-test. **(D)** *Cpeb4* mRNA levels in HuR RNA-immunoprecipitates (IP) performed in wild-type and *p38α*MKO BMDMs, stimulated with LPS when indicated. IgG IPs served as control. IP/Input enrichment is shown, normalized to wild-type IP LPS (n=2). See also Figure S3C. **(E)** *Cpeb4* and *Tnf* decay rates in wild-type and TTPMKO BMDMs stimulated for 6 h with LPS (data from(Sedlyarov et al., 2016)). See also Figure S3D. **F-G** U2OS cells were treated with tetracyline to induce the expression of a constitutively active MKK6, which induces p38 MAPK activation(Trempolec et al., 2017). **(F)** *Cpeb4* levels upon p38 activation (+MKK6) or inhibition with p38α inhibitors (SB or PH) (n=3). **(G)** *Cpeb4* levels in control or HuR-depleted U2OS cells, where the p38 MAPK has been activated (+MKK6) or inhibited (SB) (n=2). See also Figure S3F-S3G. **(A-D, F-G)** mRNA levels were quantified by RT-qPCR. *Gapdh* (a-b, g) or *18S* **(C)** were used to normalize. **(A-B, D, F-G)** Data are represented as mean ± SEM. **(A-B)** Statistics: Two-way ANOVA. **(F-G)** Statistics: One-way ANOVA.

### CPEB4 stabilizes mRNAs encoding negative feedback regulators of the LPS response

To address the functional contribution of CPEB4 to inflammation resolution, we performed RIP-seq to identify CPEB4-bound mRNAs in untreated and LPS-activated wild-type BMDMs, using *Cpeb4*KO BMDMs as controls **(Figure 4A)**. We identified 1173 and 1829 CPEB4-associated mRNAs in untreated and LPS-treated BMDMs, respectively **(Table S2)**. These included previously described CPEB4 targets such as *Txnip* or *Vegfa*, while negative controls such as *Gapdh* were not detected **(Figure 4B)**. The 3’-UTRs of the CPEB4 co-immunoprecipitated mRNAs were enriched in canonical CPE motifs (Pique et al., 2008) **(Figure 4C)** and also in CPEs containing A/G substitutions **(Figure S4A, S4B)**. These CPE variants have been shown to be specifically recognized by the *Drosophila* orthologue of CPEB2-4 (Orb2) (Stepien et al., 2016). Gene ontology analysis of CPEB4 targets indicated an enrichment of mRNAs encoding for components of the LPS-induced MAPK pathways **(Figure S4C).** These targets included mRNAs that participate in anti-inflammatory feedback loops that negatively regulate the LPS response, consistent with the phenotype observed in *Cpeb4*KO BMDMs. Thus, *Dusp1*, *Il1rn, Socs1, Socs3, Zfp36* (encoding for TTP) and *Tnfaip3* mRNAs specifically co-immunoprecipitated with CPEB4 **(Figure 4D)**. The binding of CPEB4 to *Txnip, Dusp1* and *Il1rn* mRNAs was further validated by RT-qPCR, using *Gapdh* as negative control **(Figure 4E)**. The functional importance of CPEB4 association was confirmed by the observation that *Socs1* mRNA **(Figure 4F)** and SOCS1 protein **(Figure 4G)** levels were both reduced in LPS-treated *Cpeb4*KO BMDMs compared with wild-type BMDMs. Likewise, the levels of *Dusp1*, *Il1rn, Socs3, Tnfaip3* and *Zfp36* mRNAs were similarly reduced in LPS-treated *Cpeb4*KO BMDMs **(Figure 4H)**. To examine whether these changes in mRNA levels originated transcriptionally or post-transcriptionally, we measured the stability of *Socs1* and *Il1rn* mRNAs in wild-type and *Cpeb4*KO BMDMs stimulated with LPS and treated with actinomycin D. Both *Socs1* and *Il1rn* mRNAs displayed reduced stability in the absence of CPEB4 **(Figure 4I)**. To confirm that this effect was mediated by the presence of CPEs in the mRNA 3’-UTRs, we studied the stability of reporter mRNAs with or without CPEs in LPS-stimulated RAW264 macrophages. Indeed, after 6 h of LPS treatment, the stability of a reporter with CPEs was increased compared with the same mRNAs where the CPEs were inactivated by point mutations **(Figure 4J**). Taken together, our results suggest that CPEB4 sustains the expression of anti-inflammatory factors at late times following LPS stimulation by binding to the corresponding mRNAs and promoting their stabilization.

**Figure 4.**
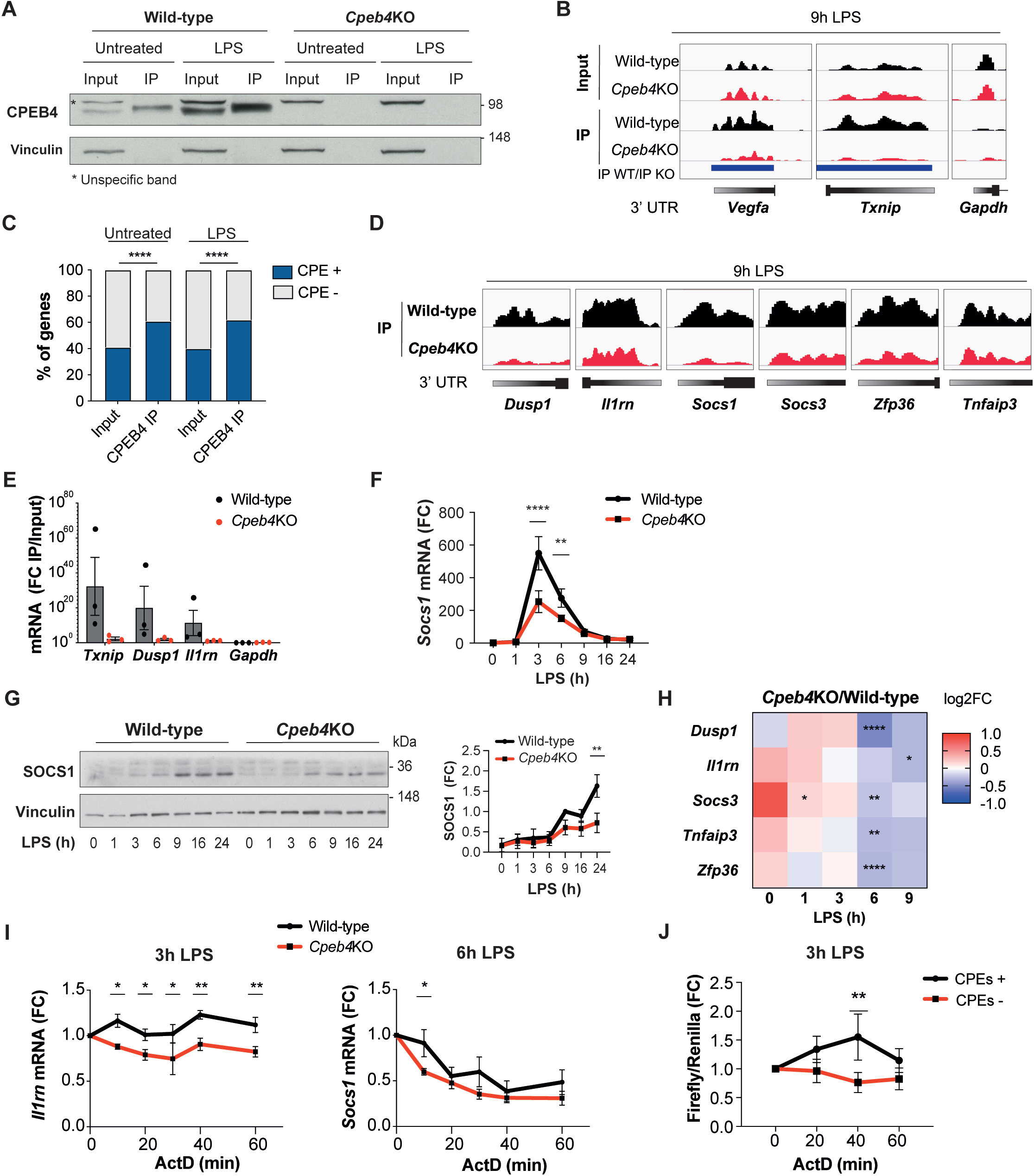
CPEB4 stabilizes mRNAs encoding negative feedback regulators of the LPS response. **A-D** CPEB4 RNA-Immunoprecipitation (IP) and sequencing was performed in total lysates (Input) from wild-type and *Cpeb4*KO BMDMs, untreated or stimulated with LPS for 9 h (n=1). **(A)** CPEB4 Immunoblot, vinculin served as a loading control. **(B)** Read coverage in Inputs or IPs of selected mRNAs. Peak enrichments between wild-type/*Cpeb4*KO IP are shown in blue. **(C)** CPE-containing transcripts [according to(Pique et al., 2008)] in Inputs and CPEB4 IPs. Statistics: Fisher’s exact test. **(D)** Read coverage in the IPs of selected mRNAs. **(E)** CPEB4 IP and RT-qPCR was performed in wild-type and *Cpeb4*KO BMDMs stimulated with LPS for 9 h. IP/Input enrichment is shown (n=3). **(F)** *Socs1* mRNA levels in LPS-stimulated wild-type and *Cpeb4*KO BMDMs. mRNA levels were measured by RT-qPCR normalizing to *Tbp* (n=6). **(G)** Immunoblot of SOCS1 in wild-type and *Cpeb4*KO BMDMs treated with LPS. Vinculin served as loading control. Quantification is shown (FC to wild-type 9 h LPS) (n=3). **(H)** Differential expression between wild-type and *Cpeb4*KO BMDMs treated with LPS measured by RNASeq (n=4). Statistics: Deseq2 R package. **(I)** mRNA stability was measured by treating with actinomycin D (ActD) wild-type and *Cpeb4*KO BMDMs stimulated with LPS for 3 h. Gene expression was analysed by RT-qPCR, normalizing to *Gapdh* (n=3). **(J)** RAW 264,7 macrophages were transfected with a Firefly luciferase reporter under the control of the cyclin B1 3’-UTR, either wild-type (CPE +) or with its CPE motifs mutated (CPE -). The same plasmid contained Renilla luciferase reporter as a control. Macrophages were stimulated with LPS for 3 h and then ActD was added. mRNA levels were measured by RT-qPCR. **(B, D)** Integrated genomic viewer (IGV) images. **(E-G)** Data are represented as mean ± SEM. **(F-G, I-J)** Statistics: Two-way ANOVA. See also Table S1 and Table S2.

### The equilibrium between CPEB4/CPEs and TTP/AREs defines mRNA oscillation patterns

Given that CPEB4/CPEs and TTP/AREs have opposite effects on mRNA stability, we explored the coexistence of both elements in the same 3’-UTR. We found that CPEB4-bound mRNAs in BMDMs were enriched in AREs **(Figure 5A, Table S3).** Conversely, 53% (102/193) of the TTP-bound mRNAs in BMDMs(Sedlyarov et al., 2016) **(Table S4)** were also co-immunoprecipitated by CPEB4 **(Figure 5B)**. Indeed, at genome-wide level, we observed a linear correlation between the number of CPEs and the number of AREs present in the same 3’-UTR **(Figure 5C, Table S3)**. To define how the coexistence of these two elements impact on mRNA stability, we compared the expression kinetics of mRNAs targeted by both CPEB4 and TTP, containing both CPEs and AREs but at different ratios, and the levels of these transcripts in LPS-activated BMDMs from the *Cpeb4*KO **(Figure 5D, Table S1)** and TTPMKO mice (Sedlyarov et al., 2016) **(Figure 5E)**. The induction of *Socs1* and *Il1rn* mRNA was reduced in *Cpeb4*KO macrophages, but the expression of *Cxcl1* or *Ptgs2* mRNA remained unaffected. Conversely, in the absence of TTP, *Socs1* and *Il1rn* mRNAs did not display major changes, while *Cxcl1* and *Ptgs2* mRNA levels increased. In a third group, mRNA levels, such as those of *Ccl2* and *Cxcl2,* were affected (in opposite directions) in both the *Cpeb4*KO and TTPMKO macrophages. Thus, despite being all bound by both proteins, some mRNAs were more dependent on CPEB4-mediated stabilization while others were more sensitive to TTP-mediated destabilization. Interestingly, the 3’-UTRs of *Socs1* and *Il1rn* mRNAs were more enriched in CPEs; those of *Cxcl1* and *Ptgs2* mRNA were enriched in AREs, while those of *Ccl2* and *Cxcl2* mRNA presented an intermediate position.

**Figure 5.**
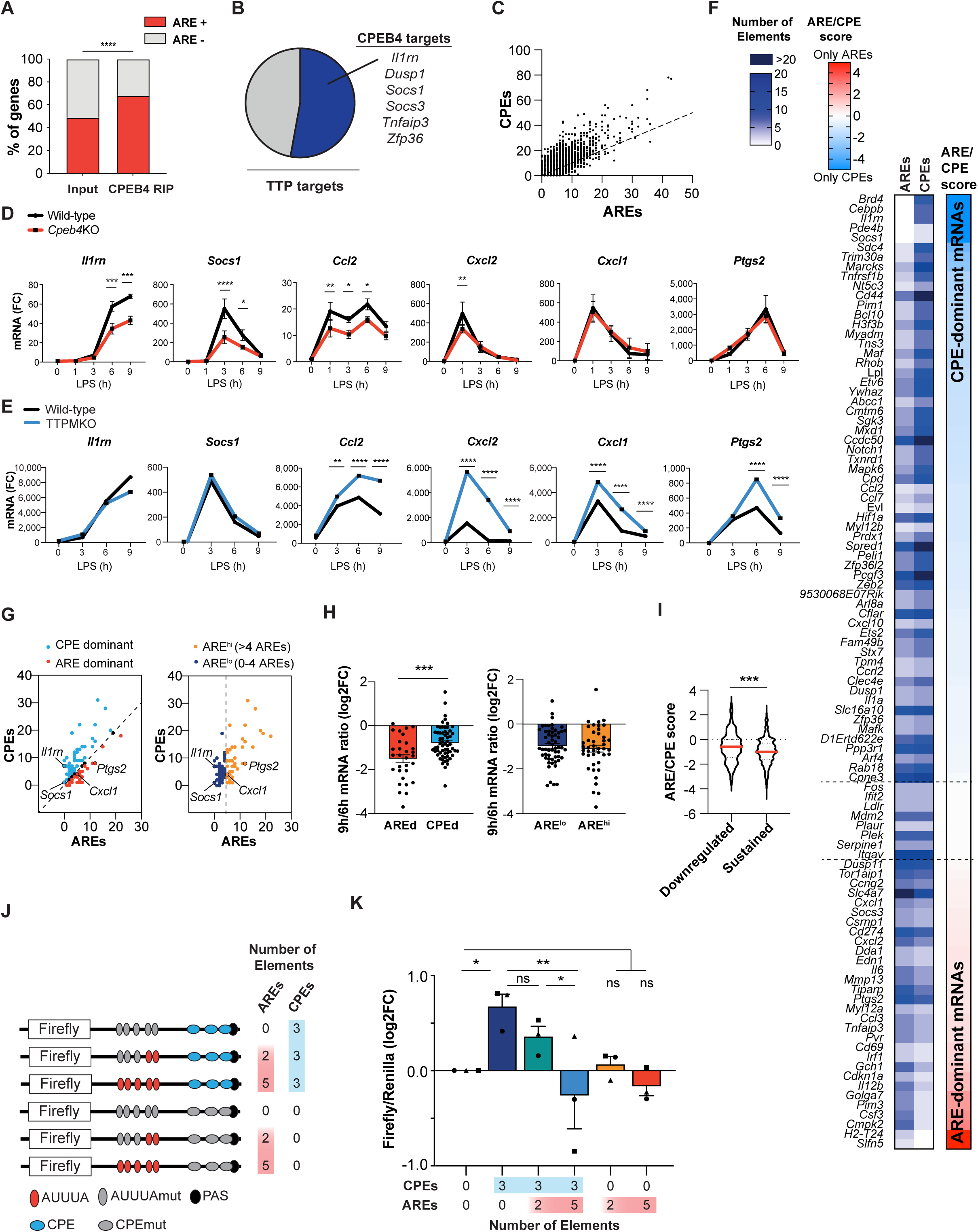
The equilibrium between CPEB4/CPEs and TTP/AREs defines different mRNA oscillation patterns. **(A)** ARE-containing transcripts in the Inputs and CPEB4 IPs from Fig. 4a-d. Statistics: Fisher’s exact test. **(B)** Overlap between CPEB4 and TTP targets(Sedlyarov et al., 2016). **(C)** Genome wide correlation between AREs and CPEs motifs in the 3’-UTRs. Black line shows the linear regression trend line. R^2^=0,6364. **(D)** mRNA levels in wild-type and *Cpeb4*KO BMDMs were measured by RT-qPCR, normalizing to *Tbp* (n=6). Statistics: Two-way ANOVA. *Socs1* data are also shown in Fig. 4f. **(E)** mRNA expression in wild-type and TTPMKO BMDMs treated with LPS. Statistics: DESeq2 software [data from(Sedlyarov et al., 2016)]. **(F)** Common TTP and CPEB4 target mRNAs were classified according to the ARE/CPE score as ARE-dominant (AREd, red) or CPE-dominant (CPEd, blue). See also Figure S5A. **(G-H)** CPEB4 and TTP target mRNAs were plotted according to the number of AREs and CPEs in the 3’-UTR. (Left) The dashed line separates AREd and CPEd mRNAs. (Right) mRNAs were classified according only to the number of AREs in their 3’UTR. The dashed lines separates ARE^high^ (>4 AREs; yellow) from ARE^low^ (*≤*4 AREs; navy) mRNAs. **H-I** Wild-type BMDMs were stimulated with LPS and mRNA levels were quantified by RNAseq (n=4). **(H)** Common CPEB4 and TTP target mRNAs were classified as AREd/CPEd (left) or ARE^high^/ARE^low^ (right). For each mRNA, the levels after 9 h of LPS treatment was normalized by its expression at 6 h LPS. **(I)** 1521 CPE-and ARE-containing mRNAs were classified as Sustained>0,5 or Downregulated<0,5 according to their expression after 9 h of LPS treatment, normalized by its peak of expression throughout LPS response. For each mRNA, its ARE/CPE score was calculated. **J-K** RAW 264,7 macrophages were transfected with a Firefly luciferase reporter under the control of a chimeric 3’-UTR combining Ier3 and Cyclin B1 AREs and CPEs motifs, respectively. Inactivating specific CPE or ARE motifs, six different 3’-UTRs with distinct ARE/CPE scores were generated. The same plasmid contained Renilla luciferase reporter as a control. **(J)** Scheme of the six constructs used for the dual luciferase reporter assay. Inactivated motifs are shown in grey. **(K)** RAW 264,7 macrophages were stimulated with LPS for 6 h and Firefly/Renilla levels were measured by RT-qPCR. Values were normalized to the 0CPEs/0AREs construct. Statistics: One-way ANOVA Friedman Test. All significant differences are shown except 5AREs/0CPEs against 3CPEs/0AREs (**) and 3CPEs/2AREs (*). **(D, H, K)** Data are represented as mean ± SEM. **(H-I)** Statistics: Mann-Whitney t test. See also Tables S2, S3, S4, S5, and S6.

Based on these observations, we classified the mRNAs that were targeted by both TTP and CPEB4 according to the ratio between the number of AREs and CPEs in their 3’-UTRs (ARE/CPE Score; **Figure 5F; Figure S5A)**. Thus, mRNAs with more CPEs than AREs were named CPE-dominant (CPEd, ARE/CPE score<0), and those with more AREs were classified as ARE-dominant (AREd, ARE/CPE score>0) **(Figure 5G; Table S3)**. In an expression kinetics analysis following LPS stimulation, we found that the decay rate at late times (6-9 h) was more pronounced for AREd than for CPEd mRNAs, while no differences were observed at early times (1-3 h) **(Figure 5H; Figure S5B, S5C; Table S5)**. When the same mRNAs were classified only as a function of their number of AREs, we found no differences in the decay rates **(Figure 5G, 5H; S5D)**. Next, we generated a list of 1521 ARE- and CPE-containing mRNAs **(Table S6)** and classified them according to their decay rate at late times after LPS stimulation, between “downregulated” (fast decay) and “sustained” (slow decay) mRNAs. Then, we interrogated if these mRNAs presented different ARE/CPE scores. We confirmed that mRNAs with slower decay rates had a lower ARE/CPE score **(Figure 5I)**. However, no differences were found when we only considered the number of AREs **(Figure S5E)**. These results indicate that the ARE/CPE ratio, but not the number of AREs alone, defines differential mRNA decay rates during the resolutive phase of the LPS response.

To directly test this model, we expressed Firefly luciferase reporter mRNAs, with different combinations of CPEs and AREs, in macrophages and measured their levels after exposure to LPS. As transfection control, we used Renilla luciferase with a control 3’-UTR (without CPEs or AREs). The parental Firefly luciferase reporter included 5 AREs and 3 CPEs, which were inactivated by point mutations to generate the following combinations: 5ARE/3CPE, 2ARE/3CPE, 0ARE/3CPE, 5ARE/0CPE, 2ARE/0CPE and 0ARE/0CPE **(Figure 5J)**. Compared with the reporter without CPEs or AREs (0ARE/0CPE), the presence of CPEs determined higher mRNA levels at late times after LPS stimulation and these levels were progressively reduced by the addition of AREs (increasing ARE/CPE score) **(Figure 5K)**. On the other hand, the number of AREs in the absence of CPEs had no significant impact on the reporter expression levels **(Figure 5K)**. No significant differences were found at early time points **(Figure S5F)**. These results confirm that the combination of CPEs and AREs on mRNAs, together with the regulation of their trans-acting factors CPEB4 and TTP, provides a temporal- and transcript-specific regulatory network that controls mRNA expression during the late phase of inflammatory processes.

## DISCUSSION

Physiological inflammatory responses rely on both rapid induction and efficient resolution to avoid the development of diseases. Differential regulation of mRNA stability plays a pivotal role in the uncoupling between the expression of pro- and anti-inflammatory mediators and is key to engage the negative feedback loops that switch-off the inflammatory response (Schott et al., 2014). Our work shows that the temporal control of inflammation is regulated by CPEB4-mediated mRNA stabilization, which acts in a coordinated manner with the well-known TTP-driven destabilization of mRNAs. The dynamic equilibrium between positive (CPEs) and negative (AREs) cis-acting elements in the mRNA 3’-UTRs, together with the regulation of their trans-acting factors CPEB4 and TTP, generates customized temporal expression profiles, which fulfil the functions needed in each phase of the inflammatory process.

Our results illustrate how CPEB4 and TTP levels and activities are integrated during the LPS response through cross-regulation at multiple post-transcriptional and post-translational layers. First, their activities are coordinately regulated by MAPK signalling pathways. Upon LPS stimulation, p38 MAPK signalling regulates TTP phosphorylation, causing a shift in the competitive binding equilibrium between HuR and TTP towards HuR, that stabilizes *Cpeb4* mRNA and promotes CPEB4 expression during the late phase of the LPS response. Then, ERK1/2 MAPK signalling (Guillén-Boixet, 2016), which is also activated by LPS, controls the progressive accumulation of the active hyperphosphorylated form of CPEB4. Moreover, CPEB4 regulates its own mRNA, generating a positive feedback loop that contributes to the increase in CPEB4 levels and activity at the late phase of the LPS response. Additionally, CPEB4 binds to the mRNA encoding TTP (*Zfp36*), adding a negative feedback loop that restores TTP activity during the late LPS response. These results underscore a superimposed layer of coordination between the MAPKs signalling pathways in the regulation of both CPEB4 and TTP levels and activities **(Figure 6A-B)**.

**Figure 6.**
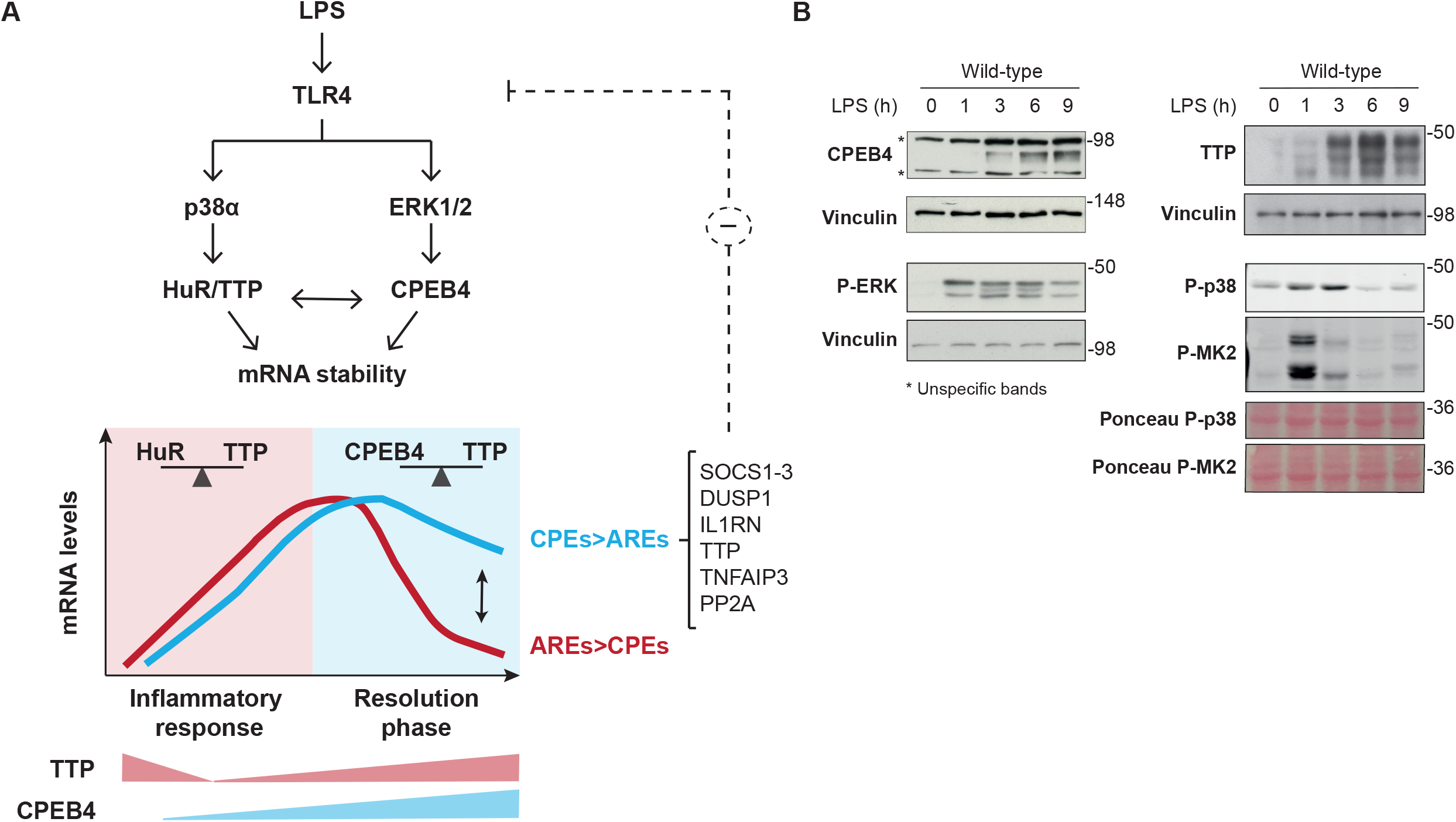
Dynamic equilibrium between TTP and CPEB4-mediated regulation of mRNAs during inflammation resolution. **(A)** LPS stimulates the MAPK signalling cascades downstream of TLR4. p38α controls TTP phosphorylation, causing a shift in the competitive binding equilibrium between HuR and TTP towards HuR, which stabilizes ARE-containing mRNAs. The p38α/HuR/TTP axis also regulates *Cpeb4* mRNA stability and promotes CPEB4 expression during the late phase of the LPS response. CPEB4 then accumulates in its active state, which involves phosphorylation by ERK1/2 MAPK signalling. During the resolutive phase of the LPS response, CPEB4 and TTP compete to stabilize/destabilize mRNAs containing CPEs and AREs. The equilibrium between these positive and negative cis-acting elements in the mRNA 3’-UTRs generates customized temporal expression profiles. CPEB4 stabilizes CPE-dominant mRNAs, which are enriched in transcripts encoding negative regulators of MAPKs that contribute to inflammation resolution. **(B)** Immunoblot of the indicated proteins in wild-type BMDMs treated with LPS for 0-9 h. Vinculin and Ponceau staining served as loading control.

The regulatory circuit that we have uncovered provides tight temporal control of the stability of TTP and CPEB4 co-regulated mRNAs, which is further delineated by the cis-acting elements in the 3’UTR **(Figure 6A-B)**. In these transcripts, the equilibrium between CPEs and AREs provides transcript-specific behaviours that contribute to uncouple the expression of pro-inflammatory mediators, which need to be rapidly silenced, from anti-inflammatory mediators, whose synthesis need to be sustained longer. Accordingly, ARE-dominant mRNAs are enriched in transcripts encoding pro-inflammatory cytokines, which are expressed during the early phase of the LPS response (Anderson, 2010; Spasic et al., 2012). In contrast, CPE-dominant mRNAs are enriched in transcripts encoding factors that contribute to inflammation resolution during the late phase of the response.

CPEB4 targets include negative regulators of MAPKs, thereby generating a negative feedback loop that limits the extent of the inflammatory response. Accordingly, we found that CPEB4 is overexpressed in patients with sepsis, and myeloid CPEB4 KO mice have decreased survival upon septic shock associated with exacerbated inflammation. Other components of this circuit can also regulate analogous inflammatory phenotypes. Thus, TTP KO mice have exacerbated LPS-induced shock (Kafasla et al., 2014), HuR KO mice display inflammatory phenotypes (Kafasla et al., 2014), and p38α deletion in macrophages reduces the LPS response and renders mice more resistant to endotoxic shock (Kang et al., 2008).

CPEBs were originally identified as a mechanism to activate maternal mRNAs, during meiotic progression in oocytes. In that scenario, a similar unidirectional and self-sustained regulatory network generated sequential waves of polyadenylation, promoting a hierarchical translation of specific subpopulations of mRNAs at each meiotic phase through the coordinated actions of CPEBs and the ARE-binding protein C3H-4 (Belloc and Mendez, 2008; Igea and Mendez, 2010; Pique et al., 2008). Our results indicate that the resolution of inflammation is controlled by an analogous regulatory network, which is coordinated through MAPK signalling pathways, the levels and activities of ARE-BP and CPEBs, and the AREs/CPEs arrangement on inflammatory and anti-inflammatory mRNAs.

## ACKNOWLEDGMENTS

We thank the Biostatistics/Bioinformatics, Histopathology, Mouse Mutant, and Functional Genomics facilities at IRB Barcelona. The Flow Cytometry Facility of the UB/PCB and the CRG Genomic Unit are also acknowledged. We thank Dr. Mercedes Fernández and members of the labs of Dr. Angel Nebreda and Dr. Raul Méndez for useful discussion. This work was supported by grants from the Spanish Ministry of Economy and Competitiveness (MINECO, BFU2017-83561-P), the *Fundación BBVA,* the *Fundación Bancaria “la Caixa”,* the *Fundació La Marató TV3* and the Scientific Foundation of the Spanish Association Against Cancer (AECC). C.S. was the recipient of an FPI-Severo Ochoa fellowship from MINECO. IRB Barcelona is the recipient of a Severo Ochoa Award of Excellence from MINECO (Government of Spain) and is supported by the CERCA Programme (Catalan Government).

## AUTHOR CONTRIBUTIONS

CS performed all the studies and contributed to experimental design, data analysis and interpretation, wrote the manuscript and assembled the figures. AS performed part of the studies, contributed to all experimental design, data analysis and interpretation and helped with manuscript preparation. RM conceived, directed and discussed the study, wrote the manuscript and obtained funding. JM contributed to *in vivo* mouse experiments. CLC performed the experiments with ARE/CPE constructs. OR analysed the sepsis datasets and the RIPseq. ID performed the genome-wide ARE/CPE analysis in the JD laboratory. AC performed the RNAseq analysis. ER contributed to the p38MKO experiments. VC contributed to the RIPseq experiments. ARN contributed to experimental design and manuscript preparation and obtained funding.

## METHODS

### Human Sepsis Microarray Analysis

Datasets used (sepsis patients/healthy subjects): GSE65682 (n=760/42), GSE57065 (n=28/25). Affymetrix U219 arrays from the GSE65682 cohort were downloaded from NCBI GEO and processed using RMA background correction and summarization with R and Bioconductor (Gentleman et al., 2004). Technical variable assessment and adjustment was performed using the Eklund metrics (Eklund and Szallasi, 2008), adjusting for RMA IQR, PM IQR, RNA Degradation and PM Median. Differentially expressed genes were determined using limma (Ritchie et al., 2015), adjusting by gender and the selected Eklund metrics, and using a |FC|>2 and Benjamini-Hochberg p-value lower than 0.05. For volcano plots, for each gene, the probeset with highest log2RMA Interquartile Range (IQR) is shown; and the y-axis represents the – log10 Benjamini-Hochberg adjusted p-value computed via limma moderated t-test. Gene expression deconvolution of blood samples in order to obtain cell proportion estimates was performed using the CellMix package (Gaujoux and Seoighe, 2013) using the DSA built-in blood signature database (Abbas et al., 2009) and the DSA algorithm with default options. The same process was done for Affymetrix U133Plus2 arrays from GSE57065 cohort. Cpeb4 mRNA expression in human immune cell lineages was obtained from Haemosphere website (Choi et al., 2019).

### Animal Studies

To generate a Myeloid-specific *Cpeb4* KO mice (*Cpeb4*MKO), conditional *Cpeb4* animals (*Cpeb4*^lox/lox(Maillo et al., 2017)^) were crossed with LysM-Cre^T/+^ (Clausen et al., 1999) transgenic animals obtained from Jackson Laboratory. Offspring was maintained in a C57BL/6J background. Routine genotyping was performed by PCR. Bone Marrow-Derived Macrophages (BMDMs) were obtained from wild-type and Myeloid-specific *p38α* KO mice (*p38α*MKO) (Youssif et al., 2018) and full CPEB4 KO mice (Maillo et al., 2017). The mice were given free access to food and water and maintained in individually ventilated cages under specific-pathogen-free conditions (unless otherwise specified). All experimental protocols were approved by the Animal Ethics Committee at the University of Barcelona.

LPS-induced Endotoxic Shock. Wild-type and Cpeb4MKO mice were injected intraperitoneally with LPS (10 mg/kg; Santa Cruz SC-3535, E. coli 0111:B4). Animals were monitored and samples were collected at the indicated times. Mice between 2 and 5 months of age were used, matched for age and sex.

Cell culture. BMDMs were isolated from the femurs of adult mice as previously described(Celada et al., 1996). Bone marrow cells were differentiated for 7 days on bacteria-grade plastic dishes (Nirco, Ref. 140298) in DMEM supplemented with 20% (vol/vol) FBS, penicillin (100 units/ml), streptomycin (100 mg/ml), L-glutamine (5 mg/ml) and 20% L-cell conditioned medium as a source of M-CSF. The medium was renewed on day 5 when BMDMs were plated for the experiment. On day 7, media were changed to DMEM containing 10% FBS, penicillin (100 units/ml), streptomycin (100 mg/ml), L-glutamine (5 mg/ml). On day 8, BMDM were primed with LPS (10 ng/ml, E. coli 0111:B4, Santa Cruz SC-3535) for the indicated time points. To inhibit p38 MAPK, BMDMs were treated with PH797804 (1 uM) for 1 h prior to LPS treatment.

Osteosarcoma cells (U2OS) were grown as in (Trempolec et al., 2017). Briefly, U2OS cells expressing the MKK6 construct were cultured in DMEM containing 10% FBS, penicillin (100 units/ml), streptomycin (100 mg/ml), L-glutamine (5 mg/ml) in the presence 4 μg/ml Blasticidin S HCl (A11139-03, Invitrogen) and 35 μg/ml Zeozin (R250-01, Invitrogen). To activate the p38 signaling pathway, cells were treated with 1 μg/ml tetracycline (Sigma 87128-25 G) or the corresponding amount of ethanol (1:1000 dilution). For the inhibition of p38α, we used 1 μM PH-797804 (Selleckchem, S2726) or 10 μM SB203580 (Axon MedChem).

### Infection with shHuR-expressing lentivirus

To generate short hairpin RNAs (shRNAs) in a lentivirus delivery system, 293 T cells (60% confluent) were transfected with 2 different shRNA-encoding DNA, pLKO.1-shHuR03 and - shHuR04 (5 μg) (IRB Barcelona Genomic Facility), VSV-G (0.5 μg) and Δ89 packaging DNA (4.5 μg) using the calcium chloride method. After overnight incubation, the media were replaced, and 48 h later, the media containing the virus were collected and passed through PVDF filters. For cell infection, the virus-containing media were diluted 1:10 in buffer containing 6 mg/l of of polybren, incubated for 10 min and then placed on the U2OS cell monolayer. The following day media were replaced, and 24 h later the cells were subjected to selection with puromycin (2 μg/ml) for 24–48 h. Selected cell populations were cultured in media with puromycin (2 μg/ml) and used for further analysis.

### RNA stability in BMDMs

Cells were stimulated with LPS (10 ng/ml, *E. coli* 0111:B4, Santa Cruz SC-3535) for the indicated times, and control cells (time 0) were collected. Fresh medium and Actinomycin D (5-10 μg/ml, Sigma Aldrich A9415) were added, and cells were collected at the indicated times. Total RNA was isolated and cDNA was synthesized to perform RT-qPCR analysis. For each timepoint, the remaining mRNA was normalized to *Gapdh* mRNA levels. The value at time 0 was set as 100%, and the percentage of remaining mRNA was calculated for the rest of timepoints.

### RNA analysis

Total RNA was either extracted by TRIzol reagent (Invitrogen), using phenol-chlorofom, or using Maxwell following the manufacturer’s instructions (Maxwell 16 LEV SimplyRNA Cells Kit, Promega). Next, 1 μg of RNA was reverse-transcribed with oligodT and random primers using RevertAid Reverse Transcriptase (ThermoFisher) or the SuperScript™ IV First-Strand Synthesis System (Thermo Fisher), following the manufacturer’s recommendations. Quantitative real-time PCR was performed in a QuantStudio 6flex (ThermoFisher) using PowerUp SYBR Green Master (ThermoFisher). All quantifications were normalized to an endogenous control (*Tbp*, *Rpl0*, *Gapdh*, *18S*). All the oligos used for RT-qPCR are listed in Table S6. Values are normalized by the median expression in untreated wild-type animals/cells.

### Cell transfection

Murine RAW 264,7 macrophages were cultured in DMEM supplemented with 10% fetal bovine serum, 2 mM L-glutamine and 1% penicillin/streptomycin at 37°C and 5% CO2. Cells were seeded in a p6 wells/plate at a concentration of 270,000 cells/well; 24 h later cells were transiently transfected with SuperFect Transfection Reagent (Qiagen Ref. 301305), following the manufacturer’s instructions, for 2.5 h and with 1.5 μg of plasmid. Cells were washed twice with PBS and allowed to rest at 37°C for 40 h. Transfected cells were then treated with LPS (10 ng/ml, *E. coli* 0111:B4, Santa Cruz SC-3535) for the indicated times. For cells used in the stability experiments, Actinomycin D (5 μg/ml, Sigma Aldrich A9415) was added for 0-60 min and cells were collected to extract total RNA. Firefly was normalized to Renilla, and time 0 of ActD was used as reference for all timepoints.

### Plasmid construction

Firefly luciferase gene with downstream Ier3 ARE fragments were purchased from GeneArt (Thermo Fisher); three fragments containing 0, 2 or 5 AREs were cloned with BamHI and HindIII restriction enzymes in pCMV-lucRenilla plasmid (Villanueva et al., 2017) upstream of the 3’-UTR of cyclin B1 mRNA. The three CPEs of cyclin B1 3’-UTR were mutated (5’-TTTTAAT-3’à5’-TTgggAT-3’; 5’-TTTTACT-3’à5’-TTggACT-3’; 5’-TTTAAT-3’à5’-TTggAAT-3’) using the QuikChange Lightning Multi Site-Directed Mutagenesis Kit (Agilent).

### Immunoblotting

Protein extracts were quantified by DC Protein assay (Bio-Rad), and equal amounts of proteins were separated by SDS-polyacrylamide gel electrophoresis. After transfer of the proteins onto a nitrocellulose membrane (GE10600001, Sigma) for 1 h at 100mV, membranes were blocked in 5% milk, and specific proteins were labeled with the following antibodies: CPEB4 (Abcam Ab83009 / clone 149C/D10, monoclonal homemade); HuR (3A2, sc-5261 Santa Cruz); HIF1a (Cayman 10006421); Phospho-p44/42 (Erk1/2) (Thr202/Try204) (Cell Signaling Clon E10, 9106); SOCS1 (Abcam Ab9870); Phospho-p38 (Thr180/Y182) (cell signaling, 9211S); p38alpha (C-20)-G (Santa Cruz, sc-535-G); Phospho-MAPKAPK2 (Thr222) (Cell Signaling, 3044 S), Tristetraprolin (TTP, D1I3T) (Cell Signaling, 71632) and Vinculin (Abcam Ab18058). Quantification was done with ImageStudioLite software, and protein expression was normalized by loading control signal.

### Lambda protein phosphatase assay (*λ*-PPase)

BMDMs were lysed in l-PPase reaction buffer (New England BioLabs, Ipswich, MA) supplemented with 0.4% NP-40 and EDTA-free protease inhibitors (Sigma-Aldrich). *λ*-PPase (New England BioLabs) reactions were performed following the manufacturer’s instructions.

### RNASeq and data analysis. Pre-processing and normalization

Reads were aligned to the mm10 genome using STAR 2.3.0e (Dobin et al., 2013) with default parameters. Counts per genomic feature were computed with the function featureCounts R (Liao et al., 2014) of the package Rsubread (Liao et al., 2013). Counts data were normalized using the rlog function of the DESeq2 R package (Love et al., 2014).

### Differential expression at gene level

Differential expression between genotype conditions (*Cpeb4*KO vs. wild-type) was performed using the DESeq2 R package (Love et al., 2014) on the original counts data. This was done independently for every LPS time point. To study expression patterns over time for wild-type samples, we first back-transformed the average (over the four samples) rlog expression to obtain normalized counts. These were further scaled by the maximum value obtained throughout the LPS response for each gene.

### Gene set analysis

Genes were annotated to Hallmark terms (Liberzon et al., 2015) using the R package *org.Hs.eg.db* Marc Carlson (2018). Gene set analysis was then performed using the regularized log transformed matrix, in which genes with low expression (genes with less than an average count of 5 reads) were filtered out. The rotation-based approach for enrichment (Wu et al., 2010) implemented in the R package limma (Ritchie et al., 2015) was used to represent the null distribution. The maxmean enrichment statistic proposed in (Efron B, 2007), under restandardization, was considered for competitive testing.

### Visualization

To obtain a measure of the pathway activity in the transcriptomic data, we summarized each Hallmark as z-score gene signature (Lee et al., 2008). To this end, rlog normalized expression values were centered and scaled gene-wise according to the mean and the standard deviation computed across samples, which were then averaged across all genes included in that Hallmark term. In addition, a global signature was computed for all the genes in the expression matrix and then used for a-priori centering of Hallmark scores. This strategy has proven useful to avoid systematic biases caused by the gene-correlation structure present in the data and to adjust for the expectation under gene randomization, i.e., the association expected for a signature whose genes have been chosen at random(Efron B, 2007; Lee et al., 2008). All statistical analyses were carried out using R and Bioconductor (Sharma et al., 2020).

RNA-immunoprecipitation (RIP). Cpeb4KO and wild-type primary BMDMs, untreated or stimulated for 9 h with LPS (10 ng/ml), were cultured in DMEM supplemented with 10% FBS, rinsed twice with 10 ml PBS, and incubated with PBS 0.5 % formaldehyde for 5 min at room temperature under constant soft agitation to crosslink RNA-binding proteins to target RNAs. The crosslinking reaction was quenched by addition of glycine to a final concentration of 0.25 M for 5 min. Cells were washed twice with 10 ml PBS, lysed with a scraper and RIPA buffer [25 mM Tris-Cl pH7.6, 1% Nonidet P-40, 1% sodium deoxycholate, 0.1% SDS, 100 mM EDTA, 150 mM NaCl, protease inhibitor cocktail, RNase inhibitors], and sonicated for 10 min at low intensity with Standard Bioruptor Diagenode. After centrifugation (10 min, max speed, 4°C), supernatants were collected, precleared, and immunoprecipitated (4 h, 4°C, on rotation) with 10 μg anti-CPEB4 antibody (Abcam), and 50 μl Dynabeads Protein A (Invitrogen). Beads were washed 4 times with cold RIPA buffer supplemented with Protease inhibitors (Sigma-Aldrich), resuspended in 100 μl Proteinase-K buffer with 70 μg Proteinase-K (Roche), and incubated for 60 min at 65°C. RNA was extracted by standard phenol-chloroform, followed by Turbo DNA-free Kit (Ambion) treatment.

Sequencing and analysis. Samples were processed at the IRB Barcelona Functional Genomics Facility following standard procedures. Libraries were sequenced by Illumina 100bp single-end and FastQ files for Input and IP samples were aligned against the mouse reference genome mm10 with Bowtie2 version 2.2.2 (Langmead and Salzberg, 2012) in local mode accepting 1 mismatch in the read seed and using default options. Alignments were sorted and indexed with sambamba v0.5.1(Tarasov et al., 2015). Putative over-amplification artifacts (duplicated reads) were assessed and removed with the same software. Coverage tracks in TDF format for IGV were made using IGVTools2 (Tarasov et al., 2015). Quality control assessment for unaligned and aligned reads was done with FastQC 0.11 (Brown et al., 2017). Further quality control steps (Gini/Lorenz/SSD IP enrichment, PCA) were performed in R with the htSeqTools package version 1.12.0 (Planet et al., 2012). rRNA contamination was assessed using the mm10 rRNA information available in the UCSC genome browser tables. A preliminary peak calling process was performed for duplicate filtered sequences with the MACS 1.4.2 software (Zhang et al., 2008) using default options to assess overall enrichment in the IP sequenced samples against their respective Input controls. The identified peaks were annotated against the mouse reference genome mm10 annotation using the ChIPpeakAnno package version 2.16 (Zhu et al., 2010). Additionally, log2RPKM quantification for mm10 UCSC 3’-UTR genomic regions (Biomart ENSEMBL archive February 2014) in all samples was obtained using custom R scripts, and potentially enriched 3’-UTRs between wild-type and Cpeb4KO IPs were identified using the enrichedRegions function from the htSeqTools package using default options and reporting Benjamini-Yekutielli adjusted p-values (pvBY). Then, for each analyzed 3’-UTR genomic region, the maximum peak score for overlapping MACS peaks was retrieved (MaxScore). 3’-UTR regions without any overlapping MACS reported peak were given a MaxScore value of 0. Finally, CPEB4-associated mRNAs were defined as those with either (I) pvBY<0.02 and log2RPKMFC>0.5; (II) pv<0.02 and log2RPKMFC>1.2 or (III) MaxScore>500 and log2RPKMFC>0.5. Gene ontology analysis was performed using the DAVID Functional Annotation Bioinformatics Microarray Analysis (Dennis et al., 2003).

### RNA-immunoprecipitation and RT-qPCR

RIP was performed as for RIP-Seq but the obtained RNA was reverse-transcribed and analyzed by RT-qPCR (see RNA analysis). The following antibodies were used: mouse monoclonal CPEB4 (ab83009 Lot. GR95787-2, Abcam); HuR antibody (3A2, sc-5261 SantaCruz); and normal mouse IgG polyclonal antibody (12-371 Sigma). Protein A/G beads were used in each case (Invitrogen).

### CPE-containing mRNAs

For the analysis of CPE-A-containing mRNAs, the script developed by Piqué et al. was run over mm10 3’-UTR reference sequences (Biomart ENSEMBL archive February 2014). mRNAs containing a putative 3’-UTR with a CPE-mediated repression and/or activation prediction were considered CPE-containing mRNAs. For CPE-G-containing mRNAs, the same script was adapted to mRNAs containing a putative 3’-UTR with the TTTTGT motif within the optimal distances to the polyadenylation signal (PAS) established by (Pique et al., 2008). As a control for the enrichment in CPE-containing mRNAs in CPEB4 target lists, the BMDM transcriptome was defined as those transcripts with > 10 reads in the corresponding wild-type input (untreated/LPS-stimulated BMDMs).

### ARE-containing mRNAs

For each gene, the reference sequence of the longest 3’-UTR was selected. The number of AREs was calculated by scanning the corresponding 3’-UTR and counting the number of occurrences of the ARE motif (ATTTA) and overlapping versions (ATTTATTTA) up to 5 overlapping motifs. Note that k overlapping motifs count as k ARE motif. ARE-containing mRNAs were considered those with 2 or more ATTTA motifs.

### ARE/CPE score

For computation of the ARE/CPE score, the reference sequence of the longest 3’-UTR per gene was selected. Similarly to the number of AREs, the number of CPEs was calculated by scanning the corresponding 3’-UTR and counting the number of occurrences of all the CPE motifs (’TTTTAT’, ‘TTTTAAT’, ‘TTTTAAAT’, ‘TTTTACT’, ‘TTTTCAT’, ‘TTTTAAGT’, ‘TTTTGT’). A version without considering the last motif yielded no significant differences. The optimal distances to the polyadenylation signal (PAS) established by (Pique et al., 2008) were not considered for this analysis.

### TTP and HuR target mRNAs

Data were obtained from TTP and HuR PAR-iCLIP experiments in LPS-stimulated BMDMs (Sedlyarov et al., 2016). For TTP, mRNAs bound after 3 h or 6 h of LPS stimulation were considered. For HuR-bound mRNAs, PAR-iCLIP data corresponded to 6 h of LPS stimulation. Only mRNAs with HuR/TTP binding in the 3’-UTR were considered.

### ARE- and CPE-containing mRNAs

The list of 1521 ARE- and CPE-containing mRNAs contained HuR, TTP or CPEB4 target mRNAs that had at least one ARE motif and one CPE motif in their 3’-UTR. mRNAs containing only AREs or only CPEs were excluded.

### mRNA levels/stability in TTPMKO BMDMs

Data were obtained from (Sedlyarov et al., 2016). The statistical analysis of the publication was also considered.

### Statistics

Data are expressed as means±SEM. Dataset statistics were analyzed using the GraphPad Prism software. For two group comparisons, column statistics were calculated and based on Standard Deviation, d’Agostino&Pearson normality test, parametric t test (assuming or not same SD) or non-parametric t-test (Mann-Whitney) was performed. For multiple comparisons, one-way ANOVA Kruskal Wallis test or two-way ANOVA followed by the Bonferroni post-hoc test was used. To assess mouse survival, Kaplan-Meier survival curves were computed with R2.15 and the survival package version 2.37-2 (Pique et al., 2008) using the likelihood-ratio test, adjusting by Gender, Weight and LPS dose. Fisher’s exact test was used for contingency analysis. Differences under p<0.05 were considered statistically significant (*pv<0.05, **pv<0.01, ***pv<0.001; ****pv<0.0001). Differences under pv<0.1 are indicated as +. For animal studies, every effort was made to include more than five mice in each group. Animals were grouped in blocks to control for experimental variability. Furthermore, wild-type and knockout animals were bred in the same cages whenever possible and the experimenter was blinded until completion of the experimental analysis. Experiments were repeated independently with similar results, as indicated in the figure legends.

### Data availability

RIP-seq and RNA-seq datasets are available in GEO (accession number GSE160191 and GSE160346, respectively). Correspondence and requests for materials or supporting raw data should be addressed to R.M.

## SUPPLEMENTAL INFORMATION

**Figure S1.**
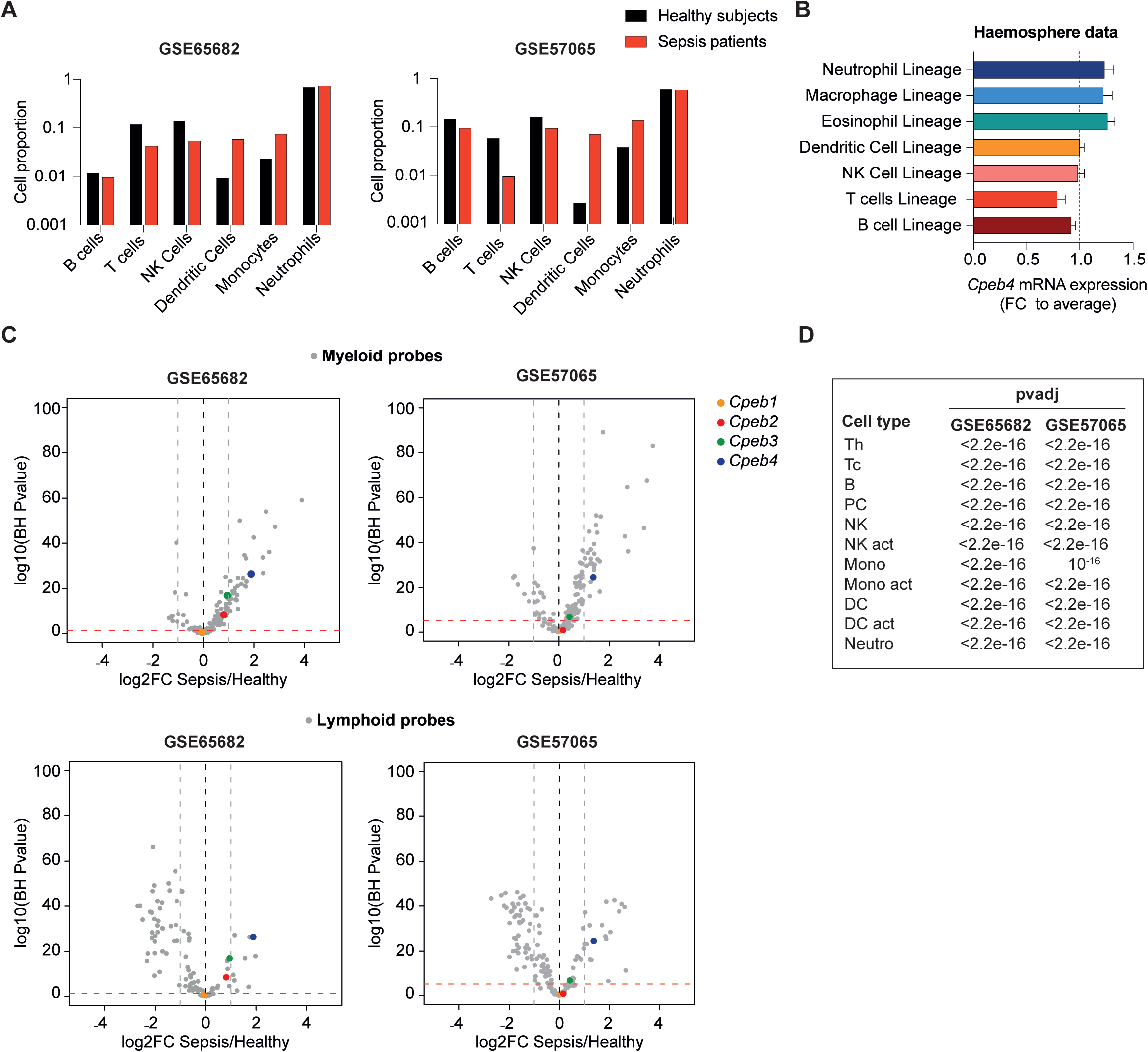
Gene expression deconvolution analysis of blood samples from septic individuals and healthy patients. **(A)** Gene expression deconvolution was performed to estimate cell proportion contribution of blood samples from septic patients and healthy subjects. **(B)** *Cpeb4* mRNA expression in human immune cell lineages (Haemosphere data). **(C)** Volcano plots showing differential expression of Smith/Abbas myeloid and lymphoid signature genes (grey) and *Cpebs* (colored) in Septic patients vs. Healthy subjects. Horizontal red dashed lines represent a pvadj of 0.05. Vertical grey dashed lines represent log2FC of -1 or 1 thresholds. **(D)** p-values for *Cpeb4* differential expression between septic patients and healthy subjects. Gene expression deconvolution was performed to adjust for the different estimated cell proportions between septic and healthy individuals.

**Figure S2.**
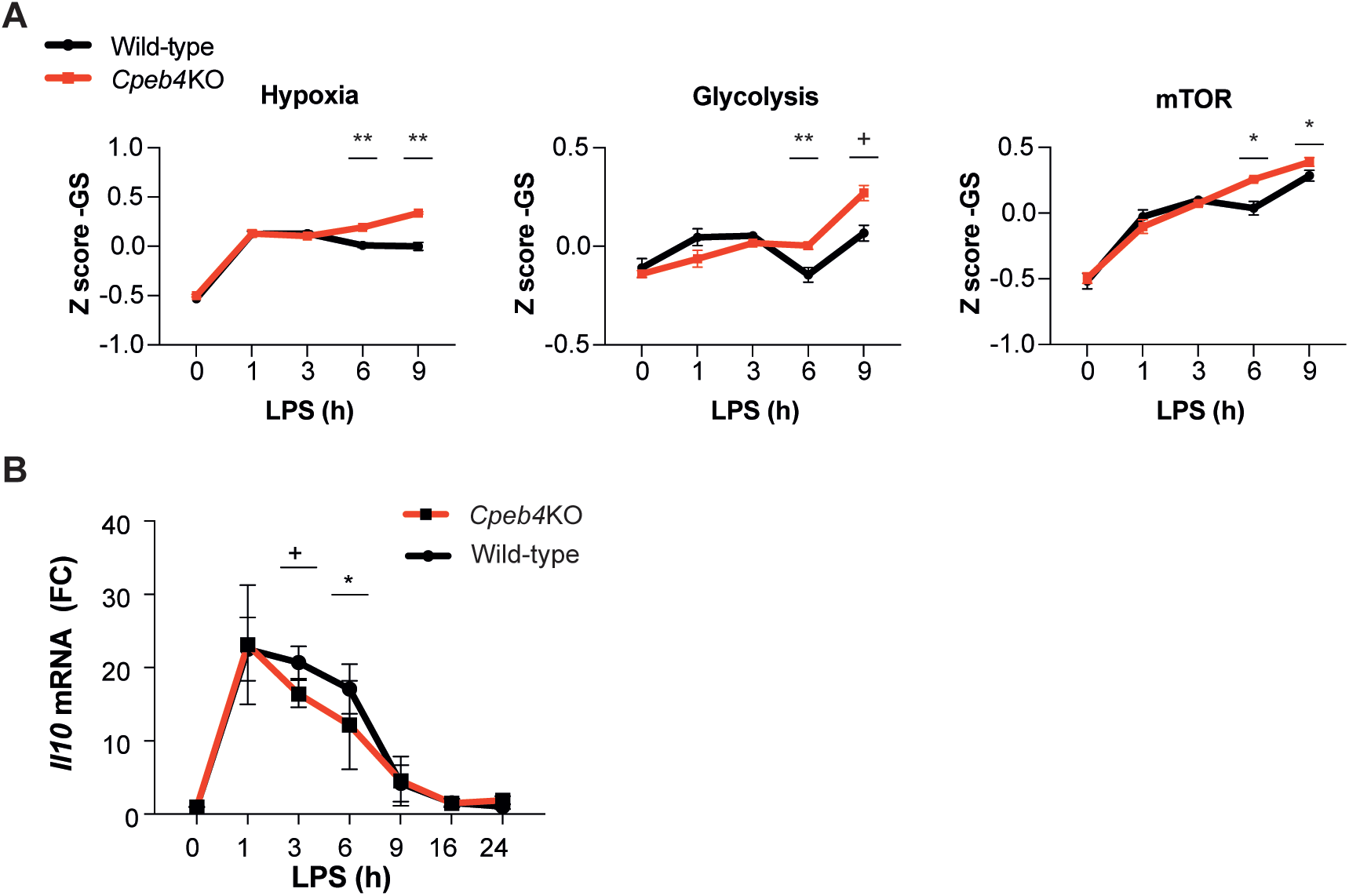
Inflammation resolution is impaired in *Cpeb4*KO macrophages. **(A)** Z-score signature of the indicated pathways in wild-type and *Cpeb4*KO BMDMs stimulated with LPS. mRNA levels were quantified by RNASeq (n=4). Statistics: rotation gene set enrichment analysis. See also Table S1. **(B)** *Il10* mRNA levels in wild-type and *Cpeb4*KO BMDMs stimulated with LPS. mRNA levels were measured by RT-qPCR, normalizing to *Tbp* (n=6). Statistics: Two-way ANOVA.

**Figure S3.**
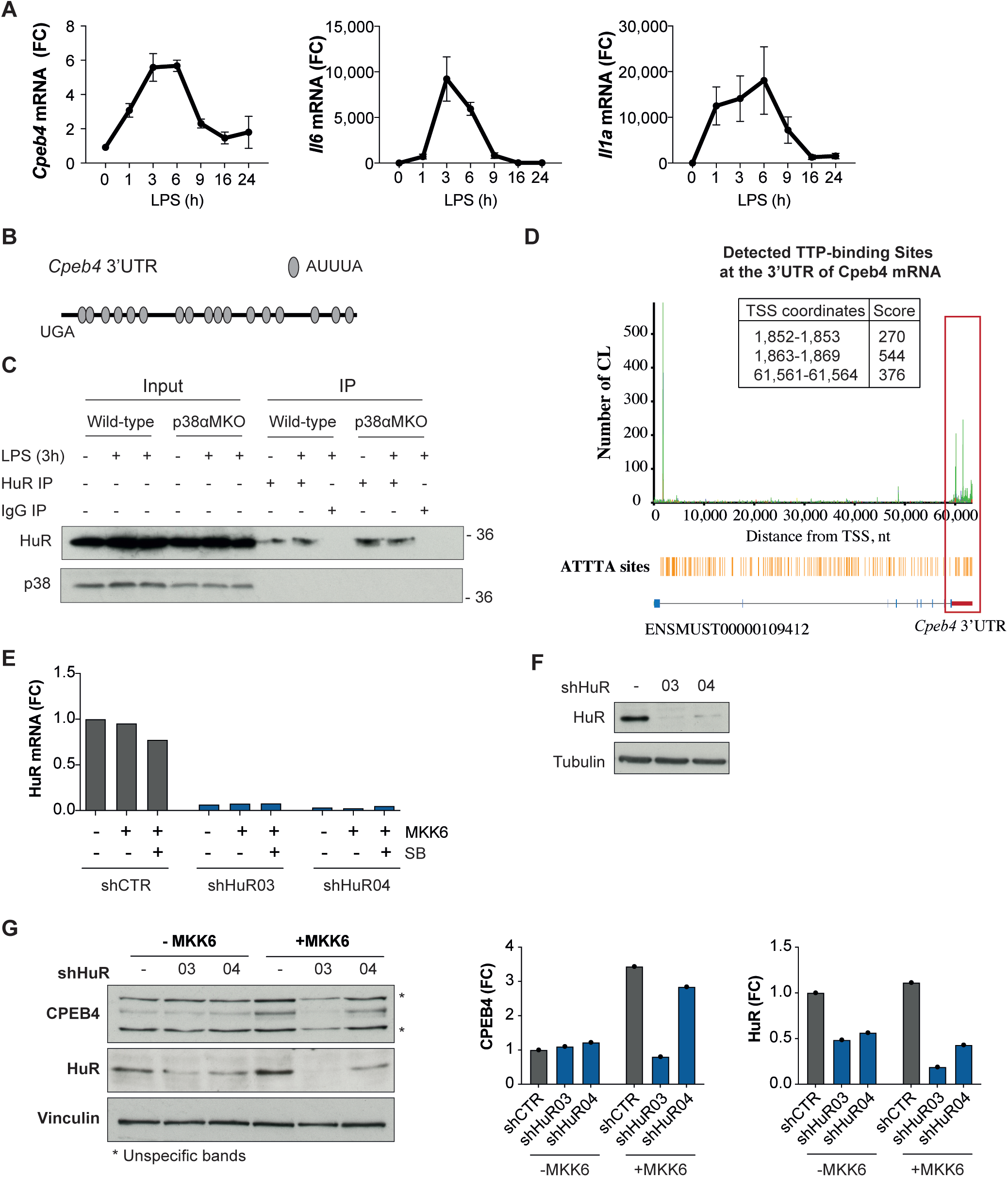
The p38*α*-HuR-TTP axis regulates *Cpeb4* mRNA stability. **(A)** BMDMs were stimulated with LPS and mRNA levels were measured by RT-qPCR, normalizing to *Tbp* (n=6). *Cpeb4* mRNA values are also shown in Fig. 2a. **(B)** Schematic representation of the *Cpeb4* 3’-UTR showing ARE domains. **(C)** HuR immunoprecipitation (IP) was performed in wild-type and p38αMKO BMDMs stimulated with LPS for 3 h when indicated. IgG IP was used as control. **(D)** TTP PAR-iCLIP was performed in BMDMs treated with LPS for 6 h. Coverage plots represent the number of crosslink sites (CL) detected in each position of *Cpeb4* mRNA. For binding sites located in the *Cpeb4* 3’UTR, distance to the Transcription Start Site (TSS) and their scores are indicated (data from Sedlyarov et al., 2016). **E-G** U2OS cells, infected with shHuR (03, 04) or shCTR (-), were treated with tetracyline to induce the expression of a constitutively active MKK6, which induces p38 MAPK activation^2^. **(E)** HuR mRNA expression was measured by RT-qPCR, normalizing to *Gapdh* (n=1). **(F)** HuR immunoblot, vinculin served as loading control. **(G)** CPEB4 and HuR immunoblot after p38 MAPK activation (+MKK6) in cells infected with shHuR or shCTR. Vinculin served as loading control. Quantification is shown.

**Figure S4.**
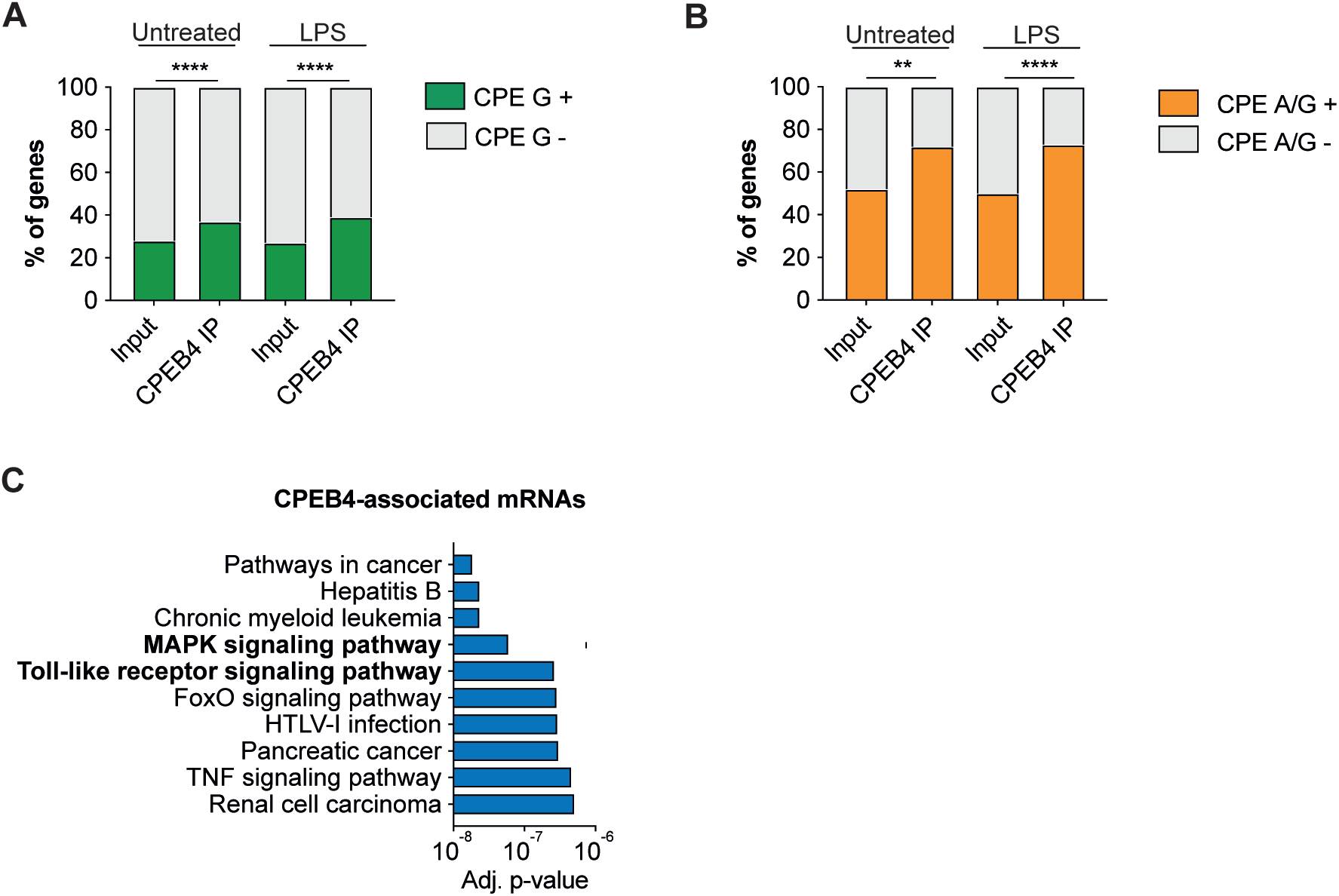
CPEB4 stabilizes mRNAs encoding negative feedback regulators of the LPS response. **A-C** CPEB4 RNA-Immunoprecipitation (IP) and sequencing was performed in total lysates (Input) from wild-type and *Cpeb4*KO BMDMs, untreated or stimulated with LPS for 9 h (n=1). **(A)** CPE-G-containing transcripts in Inputs and CPEB4 IPs. The script from Piqué et al., 2008, was modified to consider TTTTGT as a CPE motif. Statistics: Fisher’s exact test. **(B)** CPE-A- or CPE-G-containing transcripts in Inputs and CPEB4 IPs. Statistics with Fisher’s exact test. **(C)** Top 10 Gene Ontology KEGG categories enriched in CPEB4 target mRNAs in wild-type BMDMs stimulated with LPS for 9 h. *Mus musculus* transcriptome was used as background. Statistics: Benjamini-Hochberg adjusted p-value is shown. See also Table S2.

**Figure S5.**
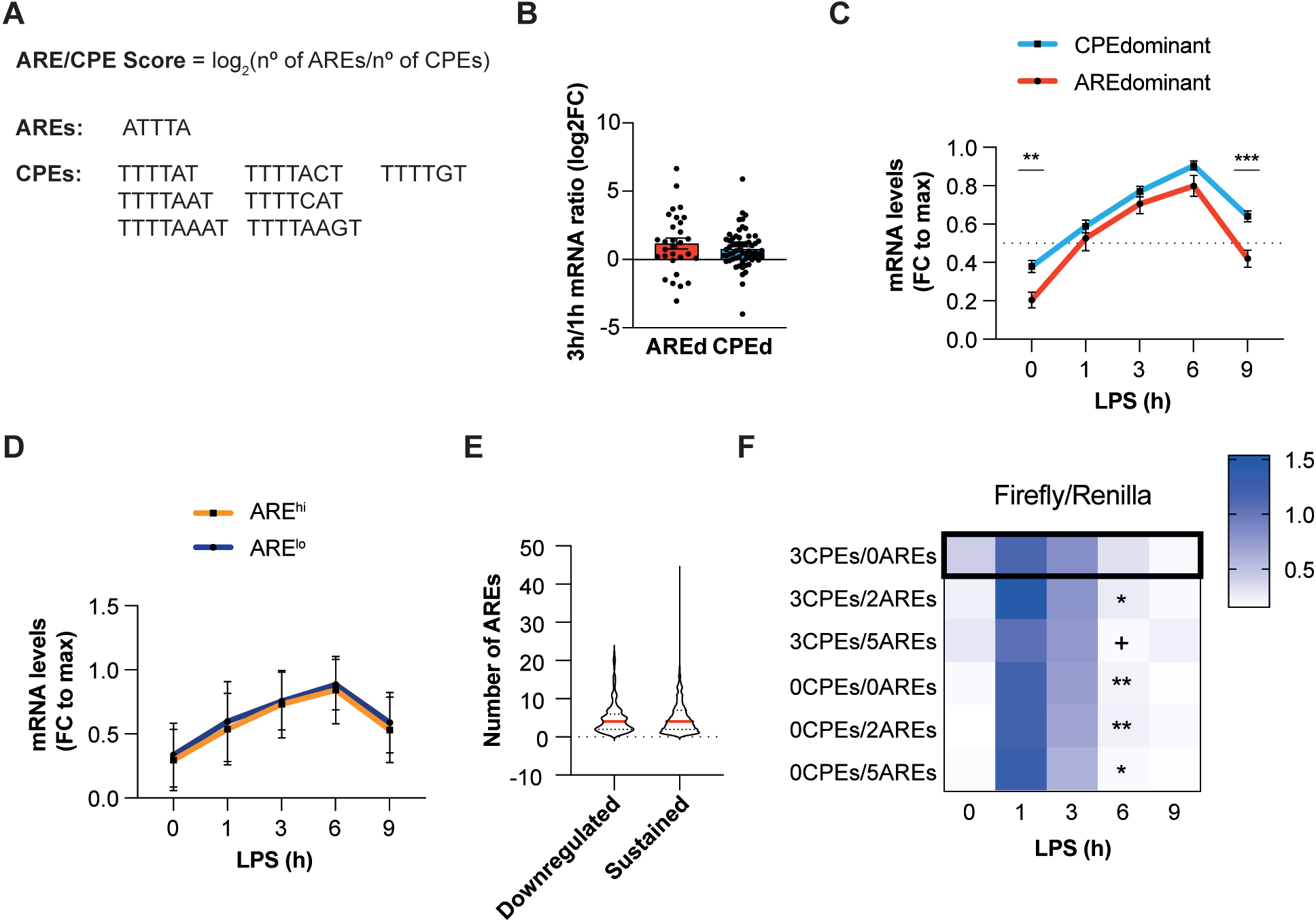
The equilibrium between CPEB4/CPEs and TTP/AREs defines mRNA oscillation patterns. **(A)** ARE/CPE score definition. ARE and CPE motifs used to calculate the score are specified. **B-E** Wild-type BMDMs were treated with LPS and mRNA levels were quantified by RNASeq (n=4). **B-D** Common TTP and CPEB4 target mRNAs were considered. **(B)** ARE-dominant (AREd, red) and CPE-dominant (CPEd, blue) mRNA levels after 3 h of LPS stimulation, normalized for its expression at 1 h. Statistics: Mann-Whitney t test. **(C)** Mean expression profile of CPEd and AREd mRNAs in LPS-stimulated BMDMs. For each mRNA, values were normalized to its peak of expression. Statistics: Two-way ANOVA. **(D)** mRNAs were plotted according to the number of AREs in the 3’-UTR as ARE^high^ (>4 AREs) or ARE^low^ (*≤*4 AREs) mRNAs. Mean mRNA expression profile of ARE^high^ and ARE^low^ mRNAs in LPS-stimulated BMDMs. For each mRNA, values were normalized to its peak of expression. **(G)** 1521 CPE- and ARE-containing mRNAs were classified as Sustained>0.5 or Downregulated<0.5, on the basis of their expression after 9 h of LPS treatment, normalized by their peak of expression during the LPS response. For each mRNA, the number of AREs in its 3’-UTR was calculated. **(H)** RAW 264,7 macrophages were transfected with a Firefly/Renilla luciferase reporter under the control of six different 3’-UTR with distinct ARE/CPE scores (See Figure 5J-K). Then, macrophages were stimulated with LPS and mRNA expression was analyzed by RT-qPCR. Heatmap shows the Firefly/Renilla expression ratio. Statistics: Two-way ANOVA vs. 3CPEs/0AREs. **(B-C, E-F)** Data are represented as mean ± SEM. See also Tables S2, S3, S4, S5, and S6.

**Table S1. RNAseq Wild-Type_vs_*Cpeb4*KO BMDMs.** Wild-type and *Cpeb4*KO BMDMs were stimulated with LPS and mRNA levels were quantified by RNASeq (n=4). Differential expression between genotype conditions was performed with Deseq2 R package. Wild-type and *Cpeb4*KO samples were compared for each time point independently.

**Table S2. RIPseq defined CPEB4 target mRNAs.** Wild-type and *Cpeb4*KO BMDMs were left untreated or stimulated with LPS for 9 h. Immunoprecipitation (IP) with anti-CPEB4 antibody was then performed, and RNA was extracted and analyzed by RNAseq. CPEB4 targets were defined based on the enrichment between wild-type and *Cpeb4*KO IPs.

**Table S3. Genome-wide AREs, CPEs and ARE/CPE Score.** For each gene, the reference sequence of the longest 3’-UTR was selected. The number of AREs and CPEs was calculated by scanning the corresponding 3’-UTR and counting the number of occurrences of each motif. The ARE/CPE score was calculated as the log2 transformed ratio between the number of ARE and CPE motifs.

**Table S4. TTP and HuR target mRNAs.** Data were obtained from TTP and HuR PAR-iCLIP experiments in LPS-stimulated BMDMs^1^. Only mRNAs with HuR/TTP binding in the 3’-UTR were considered. For TTP, mRNAs bound after 3 h or 6 h of LPS stimulation were considered. For HuR-bound mRNAs, PAR-iCLIP data corresponded to 6 h of LPS stimulation.

**Table S5. RNAseq Wild Type BMDMs.** Wild-type BMDMs were stimulated with LPS, and mRNA levels were quantified by RNASeq (n=4). Wild-type and *Cpeb4*KO samples were compared for each time point independently. The expression pattern over time for wild-type samples was analyzed (differential expression results against consecutive time points and Rlog data normalized by maximum).

**Table S6. ARE- and CPE-containing mRNAs.** ARE- and CPE-containing mRNAs were defined as all mRNAs regulated by CPEB4, TTP or HuR in LPS-stimulated BMDMs (See also Table S2 and Table S4).

**Table S7.**
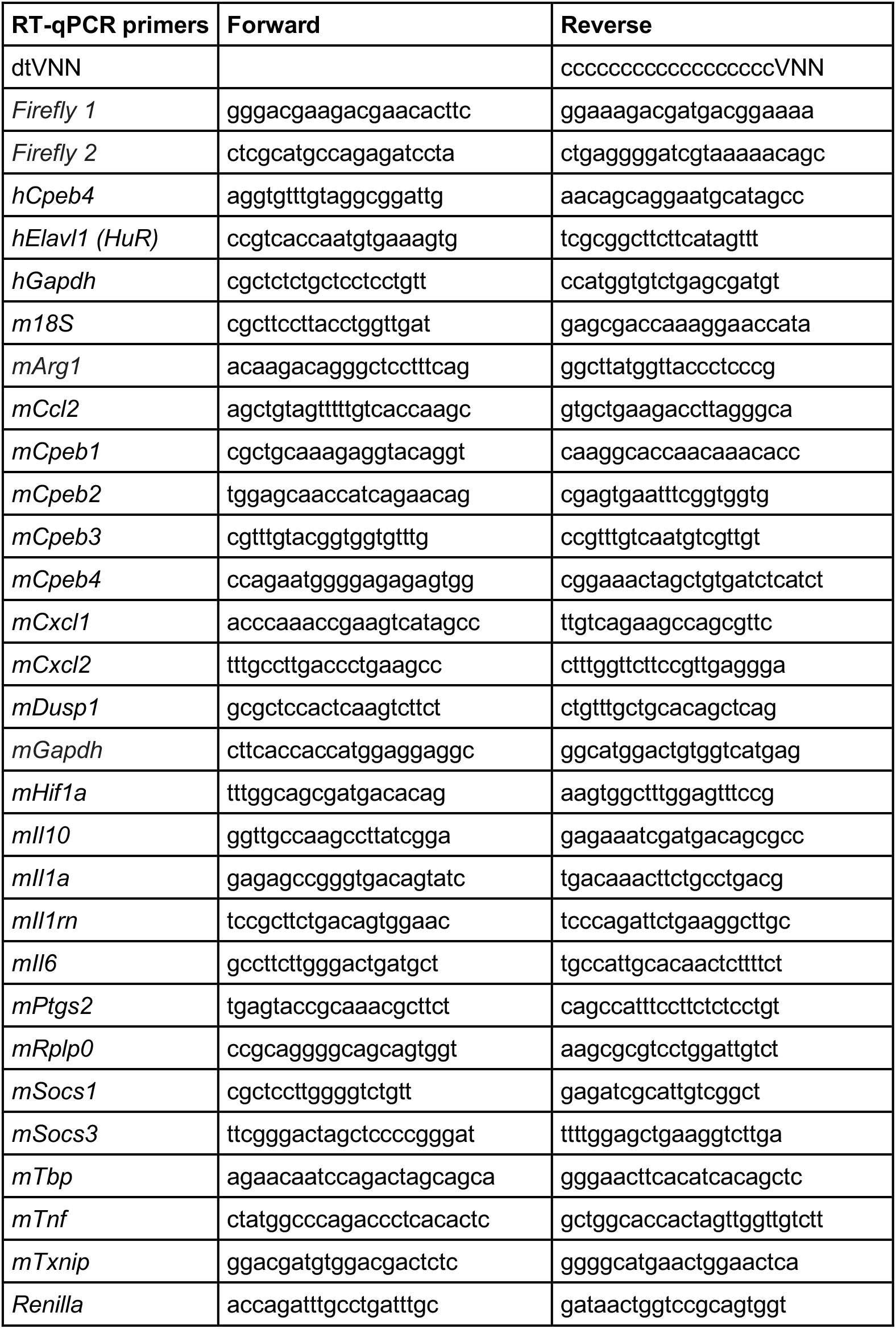
**Oligos Suñer et al.** Primers used for RT-qPCR analysis.

